# CHPT1–LCAT rewires lipolysis towards ferroptosis

**DOI:** 10.64898/2026.03.15.711301

**Authors:** Lingdi Ma, Peng Teng, Qian Zhang, Qinchi Liu, Jingjing Lu, Haoyue Wang, Yang Zhang, Zihao Guo, Ronghui Yang, Luxin Qiao, Lifang Li, Yanxia Fu, Binghui Li

## Abstract

Ferroptosis is driven by iron-dependent lipid peroxidation, yet how metabolic flux through central lipid pathways is selectively routed towards pro-ferroptotic lipid species remains unclear. Here, through an unbiased chemical–genetic screen targeting core metabolic enzymes, we identify diacylglycerol (DAG) as a licensing lipid intermediate whose pro-ferroptotic activity depends on its intracellular routing. Systematic manipulation of lipolytic flux reveals that ferroptotic vulnerability is not determined by bulk lipolytic output or downstream intermediates, but by the selective channelling of DAG into a distinct metabolic fate. Mechanistically, DAG is selectively routed through a previously unrecognized intracellular metabolic axis composed of choline phosphotransferase 1 (CHPT1) and lecithin–cholesterol acyltransferase (LCAT). We uncover an enzymatically active intracellular pool of LCAT (iLCAT) that cooperates with CHPT1 on Golgi–trans-Golgi network membranes to generate polyunsaturated cholesteryl esters that execute ferroptosis. Functionally, enforced DAG routing through this axis suppresses tumour growth via ferroptosis *in vivo*, whereas hepatocyte-specific inhibition of the CHPT1–iLCAT axis attenuates lipid peroxidation and disease progression in metabolic dysfunction–associated steatohepatitis. Together, these findings establish subcellular lipid routing, rather than lipid abundance per se, as a fundamental determinant of ferroptotic vulnerability.

---

Ferroptosis is an iron-dependent form of regulated cell death driven by lipid peroxidation and has emerged as a critical determinant of cancer progression and metabolic disease^1–4^. Although lipid metabolism is increasingly recognized as a key regulator of ferroptotic sensitivity, how specific lipid intermediates are selectively channelled into pro-ferroptotic lipid pools remains poorly understood^5,6^. In particular, whether lipid signalling intermediates merely reflect metabolic state or actively program ferroptotic vulnerability through selective subcellular routing remains unknown.

## Results

### Inhibition of MAGL sensitizes cells to ferroptosis

To systematically identify metabolic enzymes that regulate ferroptosis sensitivity, we performed a high-throughput chemical–genetic screen in HT-1080 fibrosarcoma cells, a well-established ferroptosis-sensitive model. A focused library comprising 871 small molecules targeting more than 300 metabolic enzymes involved in lipid, amino acid, nucleotide and carbohydrate metabolism was screened using a ferroptosis-rescue strategy. Cells were pretreated with either DMSO or the ferroptosis inhibitor ferrostatin-1 (Fer-1)^1^ for 12 h, followed by compound treatment for 72 h, and compounds whose cytotoxicity was selectively rescued by Fer-1 were identified (Extended Data Fig. 1a).

This screen identified multiple compounds, including the glutaminase inhibitor telaglenastat, the AMPK inhibitor compound C, and the PI3K inhibitor BEBT-908, that have previously been reported to induce ferroptosis^7–9^, thus validating the robustness of our screening platform. In addition, we uncovered five previously unreported inhibitors that target monoacylglycerol lipase (MAGL) and potently induce ferroptosis (Fig. 1a). All five compounds reduced cell viability in a dose-dependent manner, and this cytotoxicity was fully rescued by Fer-1 (Extended Data Fig. 1b), indicating that MAGL inhibition promotes ferroptosis rather than nonspecific toxicity.

**Fig. 1.**
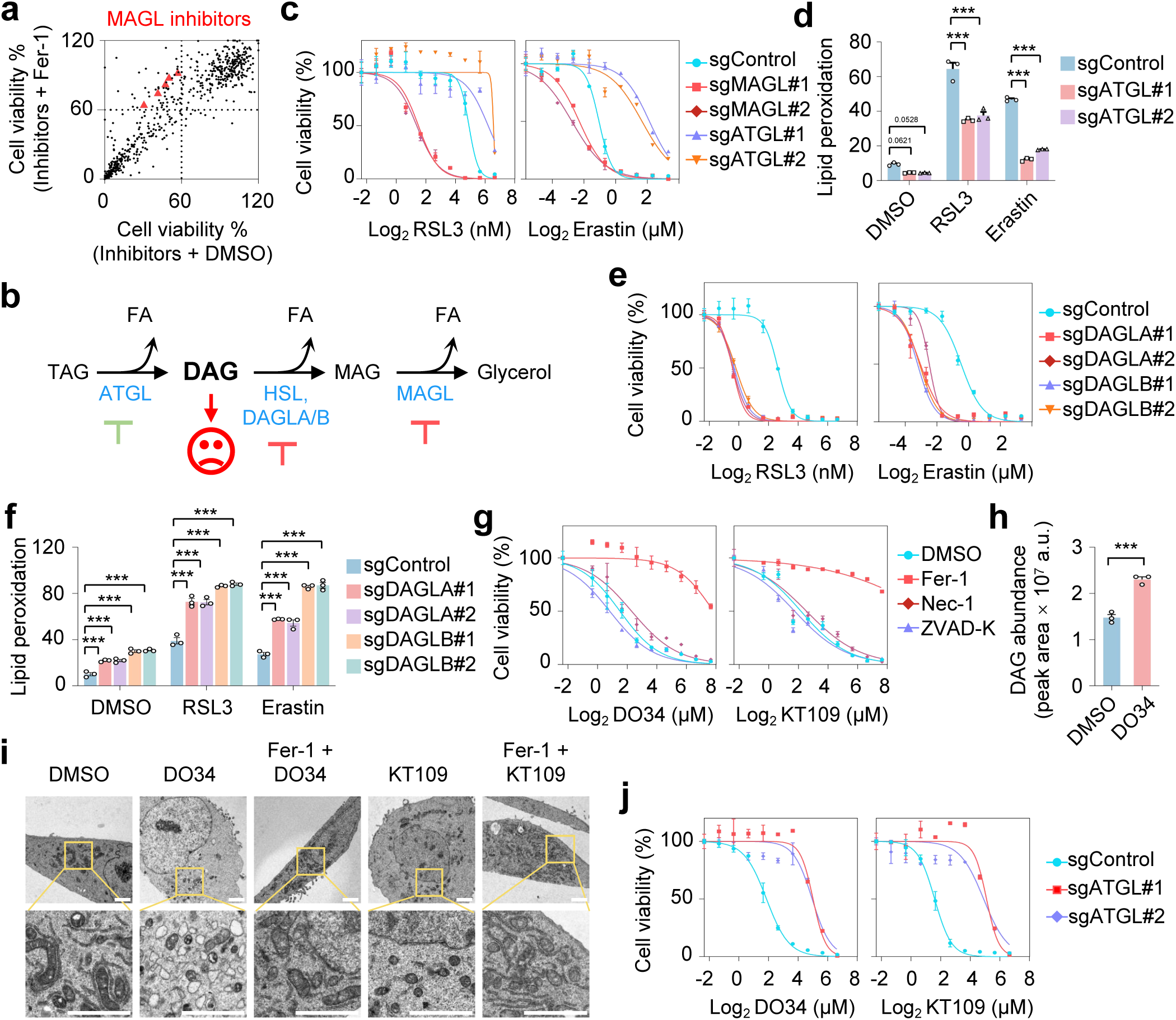
DAG is a central lipolytic node controlling ferroptosis. **a**, MAGL inhibitors identified from a viability-based chemical screen reduced HT-1080 cell viability; this effect was reversed by ferrostatin-1 (Fer-1, 5 µM). **b**, Schematic illustration of the lipolysis pathway highlighting ATGL-, DAGL- and MAGL-mediated DAG metabolism. **c**, Cell viability of HT-1080 cells with or without CRISPR–Cas9-mediated knockout of MAGL or ATGL (two independent sgRNAs each) treated with increasing concentrations of RSL3 or erastin for 24 h. **d**, Lipid peroxidation levels in ATGL-knockout HT-1080 cells treated with DMSO, RSL3 (20 nM) or erastin (10 µM) for 4 h. **e**, Cell viability of HT-1080 cells with or without DAGLA or DAGLB knockout treated with increasing concentrations of RSL3 or erastin for 24 h. **f**, Lipid peroxidation levels in HT-1080 cells with or without DAGLA or DAGLB knockout following treatment with DMSO, RSL3 (10 nM) or erastin (5 µM) for 4 h. **g**, Cell viability of HT-1080 cells treated with increasing concentrations of DO34 or KT109 for 24 h in the presence of DMSO, Fer-1 (5 µM), necrostatin-1 (Nec-1, 10 µM) or Z-VAD-FMK (Z-VAD, 10 µM). **h**, Lipidomics profiling of DAG species in HT-1080 cells after treatment with DMSO or DO34 (20 µM, 12 h). **i**, Representative transmission electron microscopy images showing mitochondrial morphology in HT-1080 cells treated with DMSO, DO34 or KT109 (20 µM) for 4 h in the presence or absence of Fer-1 (5 µM). Scale bar, 2 µm. **j**, Cell viability of ATGL-knockout HT-1080 cells treated with increasing concentrations of DO34 or KT109 for 24 h. Data in **c–h** and **j** are mean ±s.d. from three biological replicates (n = 3). Statistical analysis in **d**, **f** and **h** was performed using one-way ANOVA; *P < 0.05, **P < 0.01, ***P < 0.001.

We next focused on two well-characterized MAGL inhibitors, JW642 and JZL184^10,11^. Treatment with either compound markedly increased malondialdehyde (MDA) levels and lipid peroxidation, both of which were efficiently suppressed by Fer-1 (Extended Data Fig. 1c,d). Transmission electron microscopy (TEM) revealed hallmark mitochondrial features of ferroptosis^12^, including pronounced mitochondrial shrinkage and increased membrane density following MAGL inhibition; these changes were reversed by Fer-1 co-treatment (Extended Data Fig. 1e).

Moreover, all five MAGL inhibitors significantly sensitized cells to the canonical ferroptosis inducers RSL3 and erastin (Extended Data Fig. 1f,g). This enhanced sensitivity was fully abrogated by Fer-1 or liproxstatin-1 (Lip-1)^13,14^, but not by the apoptosis inhibitor Z-VAD-FMK^15^ or the necroptosis inhibitor necrostatin-1^16^ (Extended Data Fig. 1h,i), confirming the ferroptotic nature of cell death.

Collectively, these results identify MAGL as a negative regulator of ferroptosis, with its inhibition enhancing lipid peroxidation and ferroptotic sensitivity.

### Lipolysis-derived DAG licenses ferroptosis

MAGL catalyses the final step of lipolysis by hydrolysing monoacylglycerol (MAG), prompting us to ask whether its anti-ferroptotic function reflects altered lipolytic flux or the metabolic fate of specific lipolytic intermediates. Lipolysis is initiated at lipid droplets by ATGL-mediated hydrolysis of triacylglycerol (TAG)^17,18^ (Fig. 1b). Given previous evidence that sequestration of unsaturated lipids in lipid droplets suppresses ferroptosis^19–21^, we genetically ablated MAGL or ATGL in HT-1080 (GPX4-high) and NCI-H226 (GPX4-low) cells^22^ (Extended Data Fig. 2a,b).

Whereas MAGL deletion markedly sensitized cells to ferroptosis, loss of ATGL conferred robust resistance (Fig. 1c and Extended Data Fig. 2c). Lipid peroxidation closely mirrored these effects: MAGL ablation enhanced, whereas ATGL loss suppressed, RSL3- or erastin-induced lipid peroxidation (Fig. 1d and Extended Data Fig. 2d–f). These phenotypes were recapitulated by pharmacological inhibition of MAGL (using JW642 or JZL184) or ATGL (using atglistatin^23^) (Extended Data Fig. 2g,h).

To determine whether ferroptosis sensitivity is dictated by the build-up of a specific lipolytic intermediate or by its downstream metabolic processing, we next interrogated enzymes governing diacylglycerol (DAG) metabolism. Inhibition or knockdown of hormone-sensitive lipase (HSL) had minimal effects on ferroptosis sensitivity or lipid peroxidation (Extended Data Fig. 3a,b), possibly owing to its predominant expression in adipose tissue and its ability to hydrolyse both TAG and DAG^24^. By contrast, pharmacological inhibition of diacylglycerol lipase (DAGL)^25^, which converts DAG to MAG, potently sensitized diverse cancer cell lines to RSL3- or erastin-induced ferroptosis (Extended Data Fig. 3c–i). Concordantly, stable genetic ablation of DAGLA or DAGLB increased ferroptosis sensitivity, whereas DAGL overexpression conferred resistance to ferroptotic death (Fig. 1e,f and Extended Data Fig. 4a–h).

Among nine cancer cell lines examined, DAGL inhibition alone was sufficient to trigger ferroptosis in HT-1080 and SK-HEP-1 cells (Extended Data Fig. 5a,b), an effect reversed by Fer-1 but not by the apoptosis inhibitor Z-VAD-FMK or the necroptosis inhibitor necrostatin-1 (Fig. 1g and Extended Data Fig. 5c). DAGL inhibition induced DAG accumulation (Fig. 1h), and markedly elevated MDA levels and lipid peroxidation (Extended Data Fig. 5d–f). Consistent with ferroptotic cell death, TEM revealed pronounced mitochondrial shrinkage and increased membrane density upon DAGL inhibition, which were abolished by Fer-1 co-treatment (Fig. 1i). Importantly, ATGL inhibition or deletion effectively suppressed DAGL inhibition-induced lipid peroxidation and ferroptosis in both HT-1080 and SK-HEP-1 cells (Fig. 1j and Extended Data Fig. 4i–n).

Collectively, these findings establish DAG as a central lipolytic node controlling ferroptosis.

### Cholesteryl esters act downstream of DAG to execute ferroptosis

Although DAG can potentially activate PKC signaling, the PKC inhibitor GF109203X did not mitigate DAGL inhibition–induced lipid peroxidation or ferroptosis (Extended Data Fig. 6a,b). To investigate whether—and how—DAG is selectively routed to promote ferroptosis, we performed untargeted lipidomic profiling. Both DAGL and MAGL inhibition led to a marked increase in multiple unsaturated cholesteryl esters (ChEs), whereas ATGL inhibition produced the opposite effect (Extended Data Fig. 6c,d). Targeted lipidomic analysis further confirmed that DAGL inhibition selectively elevated a broad spectrum of unsaturated ChEs, while ATGL inhibition reduced their abundance (Fig. 2a).

**Fig. 2.**
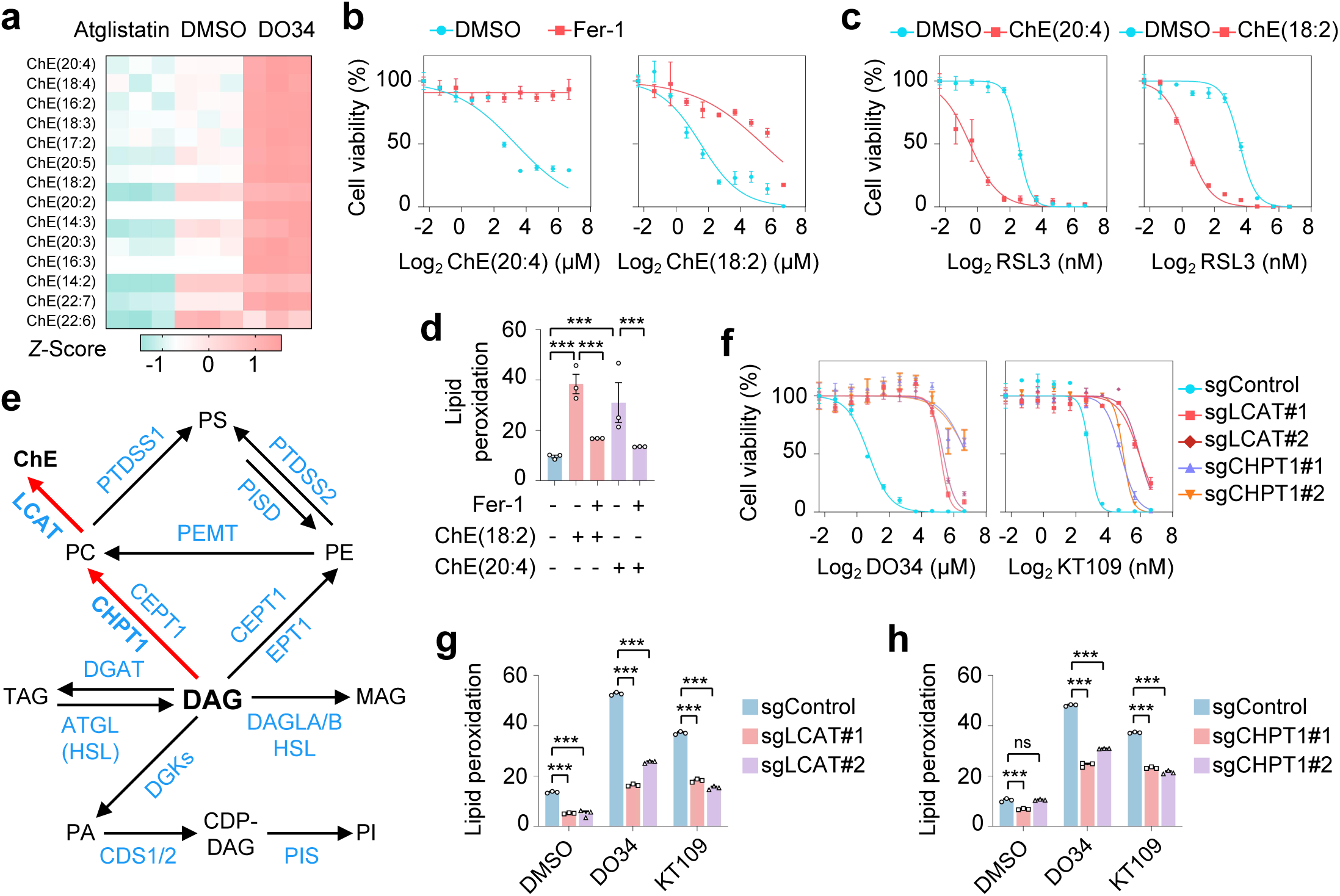
The CHPT1–LCAT axis routes DAG to promote ferroptosis. **a**, HT-1080 cells were treated with DMSO, the ATGL inhibitor atglistatin (10 µM) or DO34 (20 µM) for 4 h, followed by lipid extraction and mass spectrometry-based quantification of cholesteryl ester (ChE) species. **b**, Cell viability of HT-1080 cells treated with increasing concentrations of ChE(20:4) or ChE(18:2) for 24 h in the presence of DMSO or Fer-1 (5 µM). **c**, Cell viability of HT-1080 cells treated with increasing concentrations of RSL3 for 24 h in the presence of DMSO, ChE(20:4) or ChE(18:2) (20 µM). **d**, Lipid peroxidation levels in HT-1080 cells treated with or without ChE(20:4) or ChE(18:2) (20 µM) for 4 h in the presence or absence of Fer-1 (5 µM). **e**, Schematic illustration of DAG-associated phospholipid conversion pathways. **f**, Cell viability of HT-1080 cells with or without LCAT or CHPT1 knockout treated with increasing concentrations of DO34 or KT109 for 24 h. **g**,**h**, Lipid peroxidation levels in CHPT1- or LCAT-knockout HT-1080 cells treated with DMSO, DO34 or KT109 (20 µM) for 4 h. Data in **b–d** and **f–h** are mean ±s.d. from three biological replicates (n = 3). Statistical analysis in **d**, **g** and **h** was performed using one-way ANOVA; *P < 0.05, **P < 0.01, ***P < 0.001.

Given previous reports identifying polyunsaturated ChEs as pro-ferroptotic lipid species^26^, we selected the two most abundant and responsive species, ChE(20:4) and ChE(18:2), for functional validation. Exogenous supplementation of either ChE robustly induced ferroptosis and enhanced sensitivity to ferroptosis inducers in both HT-1080 and SK-HEP-1 cells (Fig. 2b,c and Extended Data Fig. 6e–g). Consistently, ChE treatment significantly increased lipid peroxidation and MDA levels (Fig. 2d and Extended Data Fig. 6h–j).

Notably, in both cell lines, ChE supplementation effectively overcame the ferroptosis resistance conferred by ATGL deletion (Extended Data Fig. 6k,l), indicating that cholesteryl esters function downstream of DAG routing to execute ferroptosis.

### The CHPT1–LCAT axis channels DAG into pro-ferroptotic cholesteryl esters

Sterol O-acyltransferases (SOATs), including SOAT1 and SOAT2, are the principal enzymes responsible for intracellular ChE synthesis and have also been implicated in the regulation of ferroptosis^26^. We therefore first examined whether SOATs contribute to lipid peroxidation induced by DAGL inhibition. Although knockdown of SOAT1 and SOAT2 modestly reduced basal lipid peroxidation, it failed to attenuate DAGL inhibition-induced lipid peroxidation in both HT-1080 and SK-HEP-1 cells (Extended Data Fig. 7a–c).

As DAG represents a central lipid intermediate that functions both as a signalling molecule and as a precursor for membrane phospholipid synthesis (Fig. 2e), we next sought to identify the metabolic pathway responsible for converting DAG into ChEs. To this end, we performed an siRNA-based screen targeting enzymes involved in DAG and phospholipid metabolism (Extended Data Fig. 7d). Notably, the candidate genes included lecithin–cholesterol acyltransferase (LCAT), a cholesterol–esterifying enzyme classically characterized as a secreted protein associated with high-density lipoproteins in plasma^27^.

This screen revealed that knockdown of either choline phosphotransferase 1 (CHPT1) or LCAT robustly suppressed the increase in lipid peroxidation induced by DAGL inhibition (Extended Data Fig. 7e–m). Consistent with their close functional coupling, CHPT1 catalyses the conversion of DAG to phosphatidylcholine (PC), whereas LCAT esterifies cholesterol using PC as the acyl donor, thereby generating ChEs^27,28^. These findings identify a direct metabolic route whereby DAG is channelled into ChE production through the sequential actions of CHPT1 and LCAT.

Consistent with the screening results, genetic ablation of either CHPT1 or LCAT in both HT-1080 and SK-HEP-1 cells (Extended Data Fig. 8a,b) markedly suppressed ferroptosis induced by DAGL inhibition, as well as ferroptosis triggered by the canonical inducers RSL3 and erastin (Fig. 2f–h and Extended Data Fig. 8c–i).

Together, these findings identify a direct metabolic route whereby DAG is selectively channelled into ChE production through the CHPT1–LCAT axis.

### An intracellular pool of LCAT spatially cooperates with CHPT1 to drive ferroptosis

LCAT is classically defined as a secreted enzyme that enters the endoplasmic reticulum via its signal peptide, undergoes glycosylation, and is secreted into the circulation, where it is activated by apolipoprotein A-I (ApoA-I) to catalyse cholesterol esterification^29^. To determine whether LCAT can also function intracellularly in the context of DAG-driven ferroptosis, we expressed full-length LCAT, a signal peptide-deficient LCAT variant, and a catalytically inactive mutant in HT-1080 and SK-HEP-1 cells (Extended Data Fig. 9a,b).

As expected, the signal peptide-deficient LCAT was not detectable in the culture medium and was retained intracellularly, whereas both full-length and catalytically inactive LCAT were efficiently secreted (Extended Data Fig. 9c), establishing that the signal peptide is required for LCAT secretion. Notably, LCAT recovered from conditioned media exhibited a higher degree of glycosylation than intracellular over-expressed LCAT, with the signal peptide-deficient form displaying the lowest glycosylation level, consistent with progressive glycosylation during passage through the secretory pathway (Extended Data Fig. 9c). Importantly, the Flag-tagged signal peptide-deficient LCAT migrated at a molecular weight comparable to that of endogenous LCAT and to bacterially expressed His-tagged LCAT, used here as a non-glycosylated reference (Extended Data Fig. 9c), indicating that the intracellular pool of endogenous LCAT is largely non- or minimally glycosylated.

Despite these marked differences in trafficking and glycosylation, overexpression of either full-length or signal peptide-deficient LCAT robustly enhanced ferroptosis and lipid peroxidation, particularly upon DAGL inhibition or treatment with RSL3 or erastin, whereas the catalytically inactive mutant failed to do so (Fig. 3a,b and Extended Data Fig. 9d–h), demonstrating that intracellular LCAT is sufficient to promote ferroptosis.

**Fig. 3.**
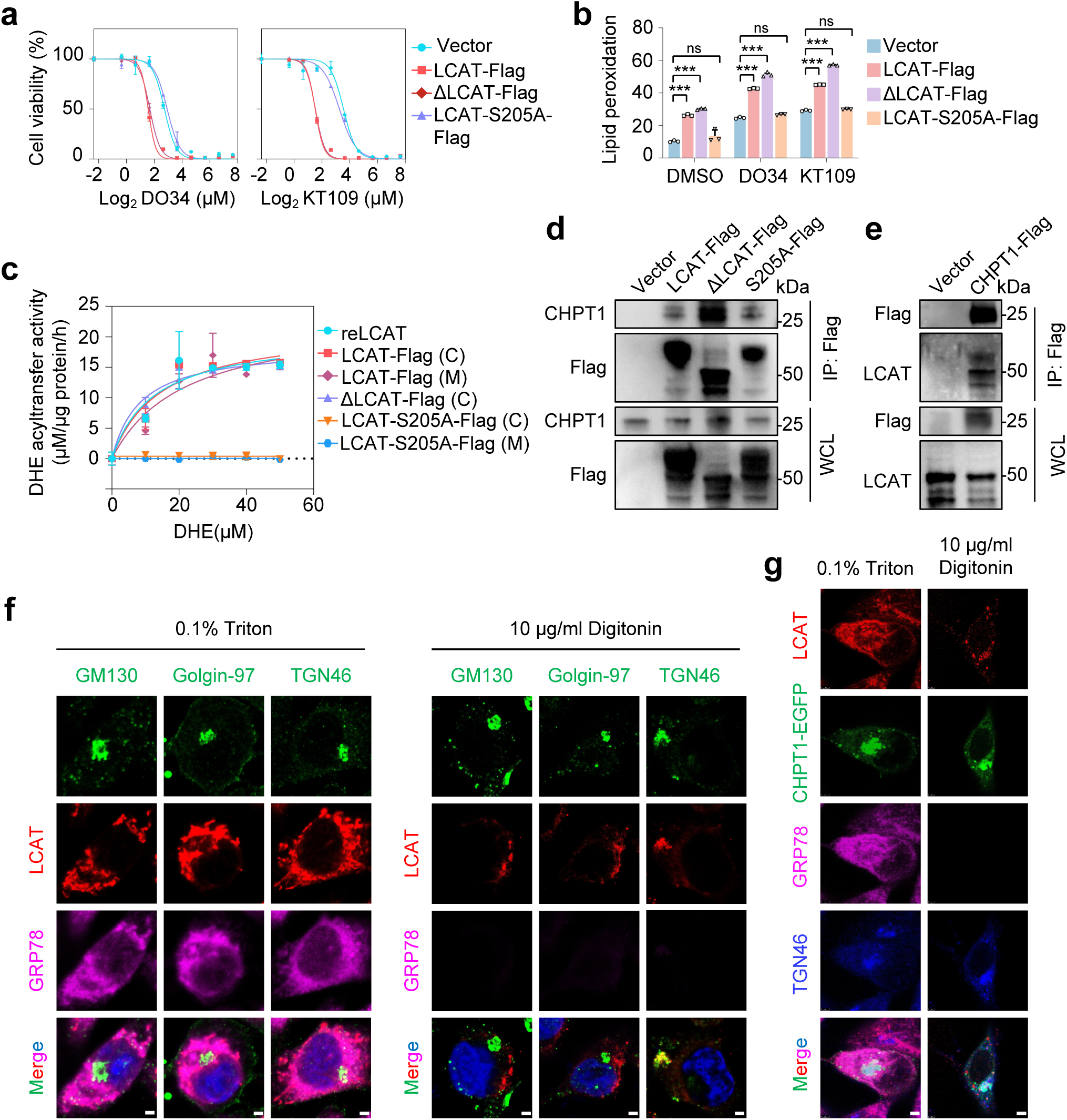
Intracellular LCAT cooperates with CHPT1 to promote ferroptosis. **a**, Cell viability of HT-1080 cells expressing empty vector, wild-type LCAT–Flag, signal peptide-deficient ΔLCAT–Flag or catalytically inactive LCAT-S205A–Flag, treated with increasing concentrations of DO34 or KT109 for 24 h. **b**, Lipid peroxidation levels in HT-1080 cells expressing the indicated LCAT–Flag constructs following treatment with DMSO, DO34 or KT109 (10 µM) for 4 h. **c**, *In vitro* DHE acyltransferase activity of purified LCAT–Flag proteins (wild-type, ΔLCAT or S205A mutant) measured across increasing DHE concentrations. reLCAT, recombinant LCAT; (C), LCAT–Flag recovered from cell lysates; (M), LCAT–Flag enriched from conditioned medium (secreted fraction). **d**, Co-immunoprecipitation of endogenous CHPT1 using the indicated LCAT–Flag constructs as bait in HEK293T cells. **e**, Co-immunoprecipitation of endogenous LCAT using CHPT1–Flag as bait in HEK293T cells. **f**, Immunofluorescence images showing localization of LCAT with Golgi markers (GM130, Golgin-97 and TGN46) and GRP78 (an endoplasmic reticulum luminal protein marker) following selective permeabilization with 0.1% Triton X-100 or 10 µg ml⁻¹digitonin. Scale bar, 2 µm. **g**, Immunofluorescence images showing co-localization of LCAT, CHPT1–GFP, GRP78 and TGN46 following selective permeabilization with 0.1% Triton X-100 or 10 µg ml⁻¹digitonin. Scale bar, 2 µm. Data in **a–c** are mean ±s.d. from three biological replicates (n = 3). Statistical analysis in **b** was performed using one-way ANOVA; ns, not significant; *P < 0.05, **P < 0.01, ***P < 0.001.

To directly assess whether intracellularly retained LCAT is catalytically competent in the absence of ApoA-I, we purified wild-type, signal peptide-deficient and catalytically inactive LCAT from cell lysates or conditioned media, as well as bacterially expressed LCAT (Extended Data Fig. 9i), and measured acyltransferase activity *in vitro*^30,31^. Intracellular full-length and truncated LCAT displayed enzymatic activity comparable to that of secreted or bacterially expressed LCAT (Fig. 3c), which was further enhanced by ApoA-I supplementation (Extended Data Fig. 9j). These findings indicate that LCAT glycosylation is dispensable for its intrinsic enzymatic activity, consistent with a previous report^32^.

Given that multiple enzymes can generate PC^33^, we next asked whether spatial organization underlies the specificity of the CHPT1–LCAT axis. Co-immunoprecipitation assays revealed that both full-length LCAT isoforms, as well as the signal peptide-deficient truncated form, interacted with endogenous CHPT1, with the signal peptide-deficient variant exhibiting a markedly stronger association (Fig. 3d). Consistently, CHPT1 robustly co-precipitated with endogenous LCAT (Fig. 3e), supporting the notion that intracellularly retained LCAT is selectively coupled to CHPT1.

We next determined the subcellular site of the interaction between intracellular CHPT1 and LCAT using differential permeabilization. Triton X-100 permeabilization revealed LCAT localization throughout the secretory pathway, whereas selective permeabilization with digitonin, which preserves organelle membranes^34^, specifically detected intracellularly retained LCAT, hereafter termed iLCAT. Under Triton X-100 conditions, LCAT predominantly localized to the endoplasmic reticulum and showed strong colocalization with the ER luminal protein GRP78^35^ (Fig. 3f). In contrast, upon digitonin permeabilization, GRP78 was not detectable owing to its luminal localization, whereas Golgi markers exposed on the cytosolic face of Golgi membranes, including GM130 (a cis-Golgi matrix protein)^36^, Golgin-97 (a trans-Golgi golgin)^37^ and TGN46 (a trans-Golgi network marker)^38^, remained readily detectable. Under these conditions, iLCAT exhibited pronounced colocalization with TGN46 (Fig. 3f).

CHPT1 localization was unaffected by the permeabilization strategy and showed colocalization with multiple Golgi markers^33,39^, including GM130, Golgin-97 and TGN46, as well as with iLCAT (Extended Data Fig. 10a). Furthermore, iLCAT colocalized with both CHPT1 and TGN46 (Fig. 3g and Extended Data Fig. 10b), indicating that CHPT1 and iLCAT converge on Golgi-associated membranes.

To assess the generality of this mechanism, we examined LCAT and CHPT1 expression across multiple cancer cell lines. Both proteins were broadly expressed, and an intracellular pool of iLCAT was consistently detected (Extended Data Fig. 11a). Knockdown of either LCAT or CHPT1 consistently suppressed DAGL inhibition-induced lipid peroxidation across diverse cell lines (Extended Data Fig. 11b–d).

Together, these findings uncover an unanticipated intracellular pool of enzymatically active LCAT that spatially cooperates with CHPT1 on Golgi-associated membranes to drive ferroptosis by spatially coupling DAG routing to ChE synthesis.

### Inhibition of DAG degradation suppresses tumour growth via ferroptosis

Recent studies have established ferroptosis induction as an effective strategy to suppress tumour growth^40^. To assess whether selective intracellular routing of DAG into pro-ferroptotic lipid pools restrains tumour progression *in vivo*, we performed xenograft experiments using HT-1080 and SK-HEP-1 cells.

In HT-1080 xenografts, pharmacological inhibition of MAGL with JW642 markedly suppressed tumour growth compared with vehicle-treated controls (Fig. 4a and Extended Data Fig. 12a,b). Notably, co-treatment with the clinically approved ferroptosis inducer sulfasalazine (SAS)^41^ further enhanced tumour suppression, indicating a synergistic anti-tumour effect.

**Fig. 4.**
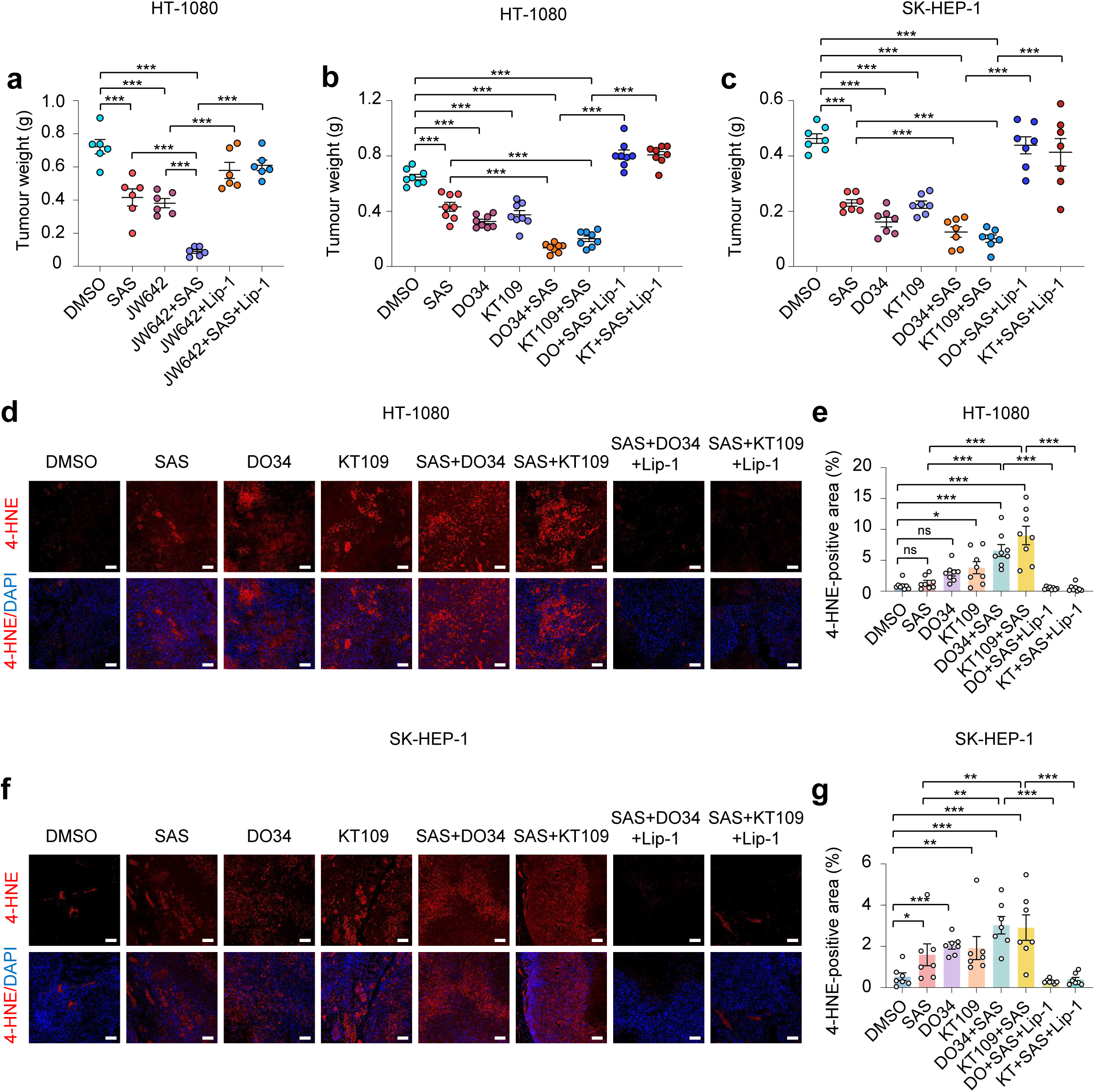
Inhibition of DAG degradation suppresses tumour growth via ferroptosis. **a–c**, Tumour weight of HT-1080 (**a**,**b**) and SK-HEP-1 (**c**) xenografts measured at the experimental endpoint following the indicated treatments. n = 6 (**a**), 8 (**b**) or 7 (**c**). **d**,**f**, Tumours from xenograft experiments were excised and sectioned for 4-HNE immunofluorescence staining; representative images are shown for HT-1080 (**d**) and SK-HEP-1 (**f**). Scale bar, 50 µm. **e**,**g**, Quantification of 4-HNE-positive areas corresponding to the images in **d** and **f**. n = 8 (**e**) or 7 (**g**), with three random fields quantified per sample. Data in **a–c**, **e** and **g** are mean ± s.d. Statistical analysis in **a–c**, **e** and **g** was performed using one-way ANOVA; ns, not significant; *P < 0.05, **P < 0.01, ***P < 0.001.

Consistently, inhibition of DAGL using two independent inhibitors, DO34 and KT109, robustly suppressed tumour growth in both HT-1080 and SK-HEP-1 xenograft models (Fig. 4b,c and Extended Data Fig. 12c–f). Co-administration of SAS further potentiated the anti-tumour efficacy of DAGL inhibition, whereas treatment with the ferroptosis inhibitor Lip-1 substantially reversed tumour suppression, indicating that the observed anti-tumour effects are ferroptosis dependent.

To directly assess lipid peroxidation in tumour tissues, immunofluorescence analysis revealed that treatment with DAGL inhibitors or SAS significantly increased 4-hydroxynonenal (4-HNE) staining, a marker of lipid peroxidation, with a pronounced synergistic increase upon combination treatment (Fig. 4d–g). This increase in 4-HNE was effectively abolished by Lip-1, further confirming ferroptosis as the underlying mechanism.

Together, these findings indicate that inhibiting DAG degradation restrains tumour progression by redirecting lipolytic intermediates toward pro-ferroptotic lipid routing and acts synergistically with ferroptosis-inducing agents.

### Targeting the CHPT1–LCAT axis ameliorates MASH by suppressing ferroptosis

Ferroptosis has been implicated as a key driver of metabolic dysfunction-associated steatohepatitis (MASH)^42^, a disease characterized by dysregulated lipid metabolism in which TAG accumulation alters both hepatic DAG availability and its intracellular routing. Given the high expression of CHPT1 and LCAT in the liver, we next investigated whether the CHPT1–LCAT axis contributes to ferroptosis-driven MASH progression *in vivo*.

To this end, we established a hepatocyte-targeted knockdown strategy using GalNAc-conjugated siRNAs^43^ to selectively silence CHPT1 or LCAT in mouse liver (Extended Data Fig. 13a,b). Six-week-old mice were fed an AMLN diet to induce MASH^44^ and, after 12 weeks, were treated with GalNAc-siRNAs targeting CHPT1 or LCAT, or a control GalNAc-siRNA (GalNAc-siControl), via tail-vein injection every four days for an additional 12 weeks (Fig. 5a and Extended Data Fig. 13b). Normal-diet-fed (ND) mice were used as controls.

**Fig. 5.**
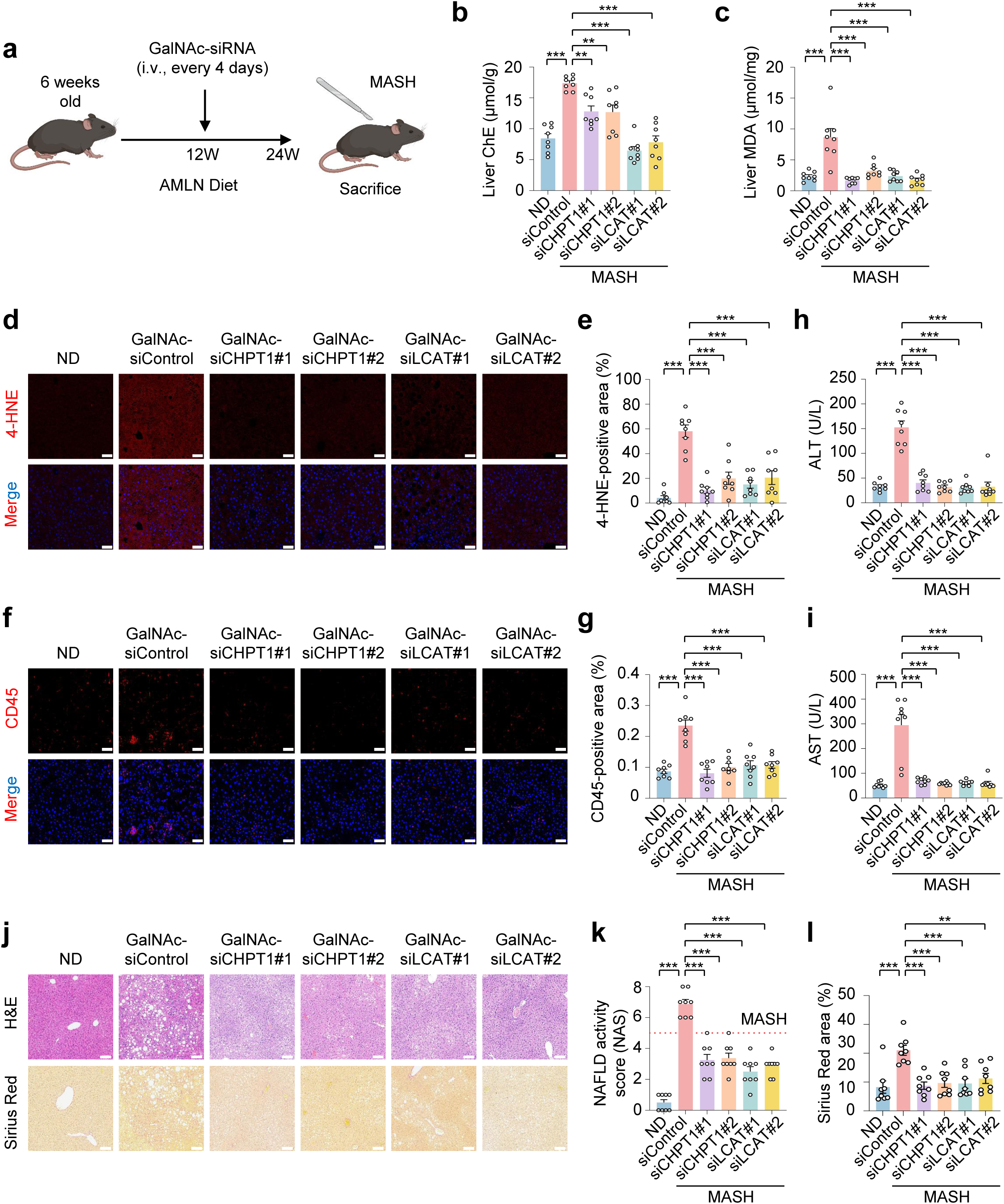
Targeting the CHPT1–LCAT axis ameliorates MASH by suppressing ferroptosis. **a**, Schematic illustration of the GalNAc–siRNA treatment strategy in an AMLN diet-induced MASH mouse model, with normal-diet-fed (ND) mice as controls. **b**,**c**, Hepatic ChE (**b**) and malondialdehyde (MDA) (**c**) levels in mouse livers at the experimental endpoint. **d**,**e**, Representative images of 4-HNE staining (**d**) in mouse liver tissues and quantification (**e**). **f**,**g**, Immunofluorescence staining of CD45 (**f**) in mouse liver sections and corresponding quantification (**g**). Scale bar, 50 µm (**d**,**f**). Quantification in **e** and **g** was performed with eight mice per group, with three randomly selected fields analysed per animal. **h**,**i**, Serum alanine aminotransferase (ALT) (**h**) and aspartate aminotransferase (AST) (**i**) levels measured at the endpoint of the MASH experiment. **j–l**, Representative haematoxylin and eosin (H&E) and Sirius Red staining of mouse liver sections (**j**) and corresponding quantitative analyses (**k**,**l**). Scale bar, 100 µm. Quantification in **l** was performed with eight mice per group, with three randomly selected fields analysed per animal. All experiments were performed with eight mice per group. Data in **b**, **c**, **e**, **g**, **h**, **i**, **k** and **l** are mean ±s.d. Statistical analysis in **b**, **c**, **e**, **g**, **h**, **i**, **k** and **l** was performed using one-way ANOVA; *P < 0.05, **P < 0.01, ***P < 0.001.

At the experimental endpoint, hepatic ChE levels were markedly reduced upon silencing of either gene, with LCAT knockdown exerting a more pronounced effect (Fig. 5b). This difference is likely attributable to the dual contribution of LCAT to both intracellular ChE synthesis and systemic cholesterol esterification. As the liver is the primary source of circulating, extracellular LCAT (hereafter referred to as eLCAT), hepatic LCAT silencing reduces not only iLCAT activity but also eLCAT-mediated ChE transport back to the liver. In contrast, CHPT1 knockdown selectively limits ChE production driven by iLCAT.

Consistent with reduced ChE accumulation, suppression of either CHPT1 or LCAT significantly decreased hepatic MDA levels and 4-HNE accumulation, indicating diminished lipid peroxidation in MASH livers (Fig. 5c–e). Concomitantly, hepatic inflammation was markedly attenuated, as evidenced by reduced immune cell infiltration and decreased CD45-positive staining (Fig. 5f,g). Serum alanine aminotransferase (ALT) and aspartate aminotransferase (AST) levels—key indicators of liver injury^45^—were also completely reduced following CHPT1 or LCAT knockdown (Fig. 5h,i).

By contrast, serum TAG and LDL cholesterol levels were not significantly altered, whereas serum total cholesterol and HDL cholesterol were modestly reduced upon LCAT knockdown (Extended Data Fig. 14a–d), consistent with previous reports describing the essential role of LCAT in HDL maturation and systemic cholesterol homeostasis^46,47^. Notably, hepatic TAG and total cholesterol content were significantly decreased in both CHPT1- and LCAT-knockdown groups (Extended Data Fig. 14e,f), likely reflecting an indirect improvement in hepatic metabolic function rather than direct effects on circulating lipid pools. These metabolic changes were accompanied by attenuation of AMLN diet-induced liver weight gain and a reduced liver-to-body-weight ratio, while adipose tissue mass remained largely unaffected (Extended Data Fig. 14g–l).

Histological analysis further demonstrated that hepatocyte-specific silencing of CHPT1 or LCAT markedly alleviated MASH pathology, as reflected by reduced NAFLD activity scores and attenuated hepatic fibrosis, as assessed by haematoxylin-eosin and Sirius Red staining, respectively (Fig. 5j–l). In contrast, suppression of neutral lipid accumulation, as assessed by Oil Red O staining, was modest (Extended Data Fig. 14m,n).

Importantly, hepatic knockdown of either CHPT1 or LCAT did not affect food or water intake but attenuated AMLN diet-induced body weight gain (Extended Data Fig. 14o), suggesting that improvement in liver function may indirectly contribute to systemic metabolic benefits.

Together, these data demonstrate that hepatocyte-specific inhibition of the CHPT1–LCAT axis alleviates MASH pathology by suppressing lipid peroxidation and ferroptosis *in vivo*.

## Discussion

Our study defines a previously unrecognized metabolic principle in which ferroptotic vulnerability is dictated not by bulk lipid abundance but by selective subcellular lipid routing. We identify a DAG–CHPT1–LCAT axis that channels lipolytic intermediates into cholesteryl esters on Golgi and trans-Golgi network membranes, thereby establishing a spatially confined platform for lipid peroxidation and ferroptotic execution (Extended Data Fig. 15).

A central finding of this work is the existence of an enzymatically active intracellular pool of LCAT that is functionally distinct from its canonical extracellular role in plasma lipoprotein metabolism. This intracellular pool, iLCAT, cooperates with CHPT1 at Golgi-associated membranes to couple phospholipid remodeling to ferroptosis, revealing an unexpected intracellular function for a protein classically viewed as exclusively secreted.

The identification of the Golgi/TGN as a site of pro-ferroptotic lipid remodeling expands the current ferroptosis framework beyond mitochondria and lipid droplets, highlighting subcellular compartmentalization as a key determinant of lipid oxidative stress. Notably, the same metabolic axis exerts divergent pathological effects depending on cellular context: in cancer cells, DAG-driven ferroptosis constrains tumour growth, whereas in hepatocytes, excessive activation of this pathway promotes lipid peroxidation, inflammation, and steatohepatitis.

These findings underscore subcellular lipid routing as a unifying mechanism linking lipid metabolism to cell fate decisions and suggest that selectively targeting intracellular lipid-remodelling nodes, such as CHPT1, may provide therapeutic leverage to modulate ferroptosis in cancer and metabolic disease while preserving essential systemic lipid functions.

## Methods

### Ethical approval

All procedures involving animals complied with all relevant ethical regulations and were approved by the Institutional Animal Care and Use Committee of Capital Medical University.

### Compounds and reagents

The metabolic enzyme inhibitor compound library was obtained from MedChemExpress (MCE, cat. no. HY-L146). Individual small-molecule inhibitors used in this study included JW642 (MCE, cat. no. HY-12332), JZL184 (MCE, cat. no. HY-15249), DO34 (MCE, cat. no. HY-117771), KT109 (MCE, cat. no. HY-18540), MAGL-IN-1 (MCE, cat. no. HY-111538), JJKK048 (MCE, cat. no. HY-108613), Elcubragis (MCE, cat. no. HY-117632) and sulfasalazine (MCE, cat. no. HY-14655). Ferroptosis-related reagents included RSL3 (MCE, cat. no. HY-100218A), erastin (MCE, cat. no. HY-15763), ferrostatin-1 (MCE, cat. no. HY-100579) and liproxstatin-1 (MCE, cat. no. HY-12726). Cell death pathway inhibitors necrostatin-1 (MCE, cat. no. HY-15760) and Z-VAD-FMK (MCE, cat. no. HY-16658B) were used where indicated.

Lipid and sterol-related reagents included arachidonoyl cholesterol (Aladdin, cat. no. C334780), cholesteryl linoleate (Aladdin, cat. no. C352059), dehydroergosterol (DHE; MCE, cat. no. HY-118667), and cholesterol oxidase (MCE, cat. no. HY-P2848). Recombinant apolipoprotein A-I (ApoA-I) was purchased from Beyotime (cat. no. P7280). Phospholipids used for reconstituted HDL preparation included DPPC (Avanti Polar Lipids, cat. no. 850355P) and POPC (Avanti Polar Lipids, cat. no. 850457P). Additional reagents included Triton™ X-100 (Sigma-Aldrich, cat. no. X100) and digitonin (MCE, cat. no. HY-N4000).

### Cell culture

HeLa, A549, HepG2, HEK293T, MCF7, HT-1080, SK-HEP-1, NCI-H226, SNU878 and U251 cell lines were obtained from the American Type Culture Collection (ATCC). HeLa, HepG2, HEK293T, MCF7, U251 and HT-1080 cells were cultured in Dulbecco’s modified Eagle’s medium (DMEM; Gibco) supplemented with 10% fetal bovine serum (FBS; Gibco) and 1% penicillin-streptomycin (Gibco, cat. no. 15140-122). A549, SNU878 and NCI-H226 cells were maintained in Roswell Park Memorial Institute (RPMI) 1640 medium (Gibco) containing 10% FBS and 1% penicillin-streptomycin. SK-HEP-1 cells were cultured in Eagle’s minimum essential medium (MEM; Gibco) supplemented with 10% FBS, 1% penicillin-streptomycin and 1% MEM non-essential amino acids (Gibco, cat. no. 11140035). All cells were maintained at 37 °C in a humidified incubator with 5% CO₂.

### Cell viability assay

Cells were seeded in triplicate into 96-well plates and treated with the indicated compounds for the specified times on the following day. Cell viability was assessed using the CellTiter-Glo Luminescent Cell Viability Assay (Promega, cat. no. G7570) according to the manufacturer’s instructions. In brief, 100 µl CellTiter-Glo reagent was added to each well, followed by incubation at room temperature for 10 min on a shaker. Luminescence was recorded using a TriStar² LB 942 multimode microplate reader (Berthold Technologies).

### Immunoblotting

Cells were washed twice with ice-cold phosphate-buffered saline (PBS) and lysed in pre-chilled lysis buffer (50 mM Tris-HCl, pH 7.4, 150 mM NaCl, 1 mM EDTA, 1% Nonidet P-40, 1 µg ml⁻¹ aprotinin, 1 µg ml⁻¹ leupeptin, 1 µg ml⁻¹ pepstatin, 1 mM Na₃VO₄ and 1 mM phenylmethylsulfonyl fluoride) for 15 min on ice. Protein concentrations were determined using the bicinchoninic acid (BCA) assay kit (Beyotime Biotechnology). Equal amounts of protein were resolved by SDS-PAGE and transferred onto polyvinylidene fluoride (PVDF) membranes (Millipore). Membranes were incubated with the indicated primary antibodies and detected using a chemiluminescent imaging system (SAGECREATION).

The following primary antibodies were used: anti-ATGL (Proteintech, cat. no. 55190-1-AP; 1:1,000), anti-MAGL (Proteintech, cat. no. 67846-1-Ig; 1:1,000), anti-GAPDH (Proteintech, cat. no. 60004-1-Ig; 1:10,000), anti-Lamin B1 (Proteintech, cat. no. 12987-1-AP; 1:10,000), anti-LCAT (Proteintech, cat. no. 12243-1-AP; 1:1,000), anti-CHPT1 (Santa Cruz Biotechnology, cat. no. sc-515577; 1:500), anti-DAGLB (Santa Cruz Biotechnology, cat. no. sc-514738; 1:500), anti-DAGLA (Abcam, cat. no. ab81984; 1:500), and anti-Flag (Thermo Fisher Scientific, cat. no. F1804; 1:10,000).

The following secondary antibodies were used: anti-mouse IgG (H+L)/HRP (Jackson ImmunoResearch, cat. no. 115-035-003; 1:100,000) and anti-rabbit IgG (H+L)/HRP (Jackson ImmunoResearch, cat. no. 111-035-003; 1:100,000).

### Targeted lipidomic analysis of cholesteryl esters

Cells were treated with the indicated compounds for the specified times, harvested by trypsinization, and collected into 1.5 ml microcentrifuge tubes. Samples were centrifuged at 800 rpm for 5 min at 4 °C, and the supernatants were removed. Cell pellets were subjected to lipid extraction by sequential addition of 300 µl methanol, 1 ml methyl tert-butyl ether (MTBE), and 250 µl water, with vortexing between each step. Phase separation was allowed to proceed at room temperature, and the extraction was repeated three times to ensure thorough recovery. The upper organic phases were pooled, centrifuged at 8,000 rpm for 15 min, and the supernatants were collected. Organic solvents were evaporated to dryness under a gentle stream of nitrogen, yielding a thin transparent film for subsequent analysis.

Metabolites were analysed using an LC-MS/MS system consisting of a 6500 plus QTrap mass spectrometer (AB SCIEX, USA) coupled to an ACQUITY UPLC H-Class system (Waters, USA). Chromatographic separation was performed on an ACQUITY UPLC BEH C18 column (100 × 2.1 mm, 1.7 µm; Waters) maintained at 45 °C. The mobile phases consisted of solvent A (methanol containing 0.1% formic acid and 10 mM ammonium acetate) and solvent B (acetonitrile/isopropanol, 2:5, v/v, containing 0.1% formic acid and 10 mM ammonium acetate). The flow rate was set to 0.5 ml min⁻¹ with the following gradient: 40% B for 1 min, increased linearly to 95% B over 6.5 min, further increased to 98% B over 1.5 min, and returned to 40% B at 8.1 min, with a total runtime of 10 min per sample.

Data were acquired in positive ion mode using multiple reaction monitoring (MRM). Ion transitions were optimized using authentic chemical standards. Source parameters were set as follows: nebulizer gas (Gas1), 60 psi; heater gas (Gas2), 50 psi; curtain gas, 30 psi; ion spray voltage, 5,000 V; and probe temperature, 250 °C. Data acquisition and peak integration were performed using SCIEX OS 1.6 software.

### Non-targeted lipidomics

Cells were treated with the indicated compounds for the specified times, harvested by trypsinization, and collected into 1.5 ml microcentrifuge tubes. Samples were centrifuged at 800 rpm for 5 min at 4 °C, and the supernatants were discarded. Cell pellets were subjected to lipid extraction by sequential addition of 300 µl methanol, 1 ml methyl tert-butyl ether (MTBE), and 250 µl water, with vortexing between each step. Phase separation was allowed to proceed at room temperature, and the extraction was repeated three times to ensure thorough lipid recovery. The upper organic phases were pooled, centrifuged at 8,000 rpm for 15 min, and the supernatants were collected. Organic solvents were evaporated to dryness under a gentle stream of nitrogen, yielding a thin transparent film for subsequent analysis.

Reverse-phase liquid chromatography was performed using a CORTECS C18 column (2.1 × 100 mm, 2.7 µm; Waters). Mobile phase A consisted of water/acetonitrile (40:60, v/v) containing ammonium acetate (0.77 g l⁻¹), and mobile phase B consisted of acetonitrile/isopropanol (10:90, v/v). The flow rate was set to 0.28 ml min⁻¹, and the column oven temperature was maintained at 40 °C. The gradient program was as follows: 30% B at 0–3.0 min, 33% B at 4.5 min, 45% B at 7.0 min, 52% B at 8.0 min, 58% B at 11.0 min, 66% B at 14.0 min, 70% B at 17.0 min, 75% B at 21.0 min, 98% B at 23.0–30.0 min, followed by re-equilibration to 30% B at 30.5–35.0 min.

Mass spectrometry was performed on an Orbitrap Exploris 240 mass spectrometer (Thermo Fisher Scientific) coupled to a Vanquish UHPLC system. Data were acquired in both positive and negative ion modes with spray voltages of 3.2 kV and 2.8 kV, respectively. The capillary temperature was set to 320 °C, with sheath gas and auxiliary gas flow rates of 35 and 10 (arbitrary units), respectively. Full MS scans were acquired over an m/z range of 240–2,000 in positive mode and 200–2,000 in negative mode at a resolution of 60,000. Data-dependent MS/MS spectra were acquired at a resolution of 15,000 with normalized collision energies of 15, 30 and 45 and a duty cycle of 1 s.

Lipid identification and quantification were performed using LipidSearch software (version 4.2; Thermo Fisher Scientific), which enables lipid annotation based on accurate mass and MS/MS fragmentation matching against an extensive lipid database.

### Lentiviral-mediated generation of gene knockout and overexpression cell lines

Gene knockout cell lines were generated using a lentiviral CRISPR-Cas9 system. Lentiviral particles were produced in HEK293T cells by co-transfecting lentiCRISPRv2 vectors encoding sgRNAs targeting the indicated genes together with the packaging plasmids pCMV-Dr8.91 and pCMV-VSV-G using polyethylenimine (Polysciences, cat. no. 23966-1). For overexpression constructs, pCDH-CMV-DAGLA or pCDH-CMV-DAGLB plasmids were transfected in parallel where indicated. Viral supernatants were collected 48 h after transfection and used to transduce HT-1080 and SK-HEP-1 cells in the presence of hexadimethrine bromide (10 µg ml⁻¹; Sigma-Aldrich, cat. no. H9268). Forty-eight hours after infection, cells were selected with puromycin to obtain pooled stable populations. Knockout efficiency was confirmed by immunoblotting before cells were used for subsequent cellular assays.

The sgRNA sequences used in this study were as follows: sgMAGL#1, ACACATC CCTGACGAAAACG; sgMAGL#2, AGTACCTCTTCTGCAGGTAC; sgATGL#1, CGTTCCGCTGCAGAAAGCGC; sgATGL#2, GCTGCGAGAGATGTGCAAGC; sgDAGLA#1, CCGTGGGGTCGAAGACGCAG; sgDAGLA#2, ACCAGCCATGCC CGGGATCG; sgDAGLB#1, GATCGTCCTCATGATTCTCC; sgDAGLB#2, CATCT ACAGAAACCCCCTCA; sgLCAT#1, GCGGGGGGAAGAGCACATTG; sgLCAT#2, TGGGGGCTTACCGAGGATGA; sgCHPT1#1, GGAACAAGAGTTTGTTCTTC; sgCHPT1#2, TCTTTCAGTATTTATGGCAG.

### siRNA-mediated gene silencing

Gene knockdown was achieved using small interfering RNAs (siRNAs). Cells were seeded in antibiotic-free medium one day before transfection to reach 30–50% confluence at the time of transfection. Gene-specific siRNAs or a non-targeting control siRNA (siControl) were transfected using Lipofectamine RNAiMAX (Invitrogen) according to the manufacturer’s instructions. Briefly, siRNAs were diluted in Opti-MEM reduced-serum medium and mixed with RNAiMAX for 10–20 min at room temperature to allow complex formation, followed by addition to cells at a final siRNA concentration of 50 nM. For each target gene, three independent siRNA oligonucleotides were pooled in equal amounts and used for transfection. After 6 h, the medium was replaced with fresh complete growth medium. Knockdown efficiency was assessed 48–72 h after transfection by quantitative PCR (qPCR), and cells were subsequently used for downstream functional assays.

The sequences of siRNAs used in this study are listed below.

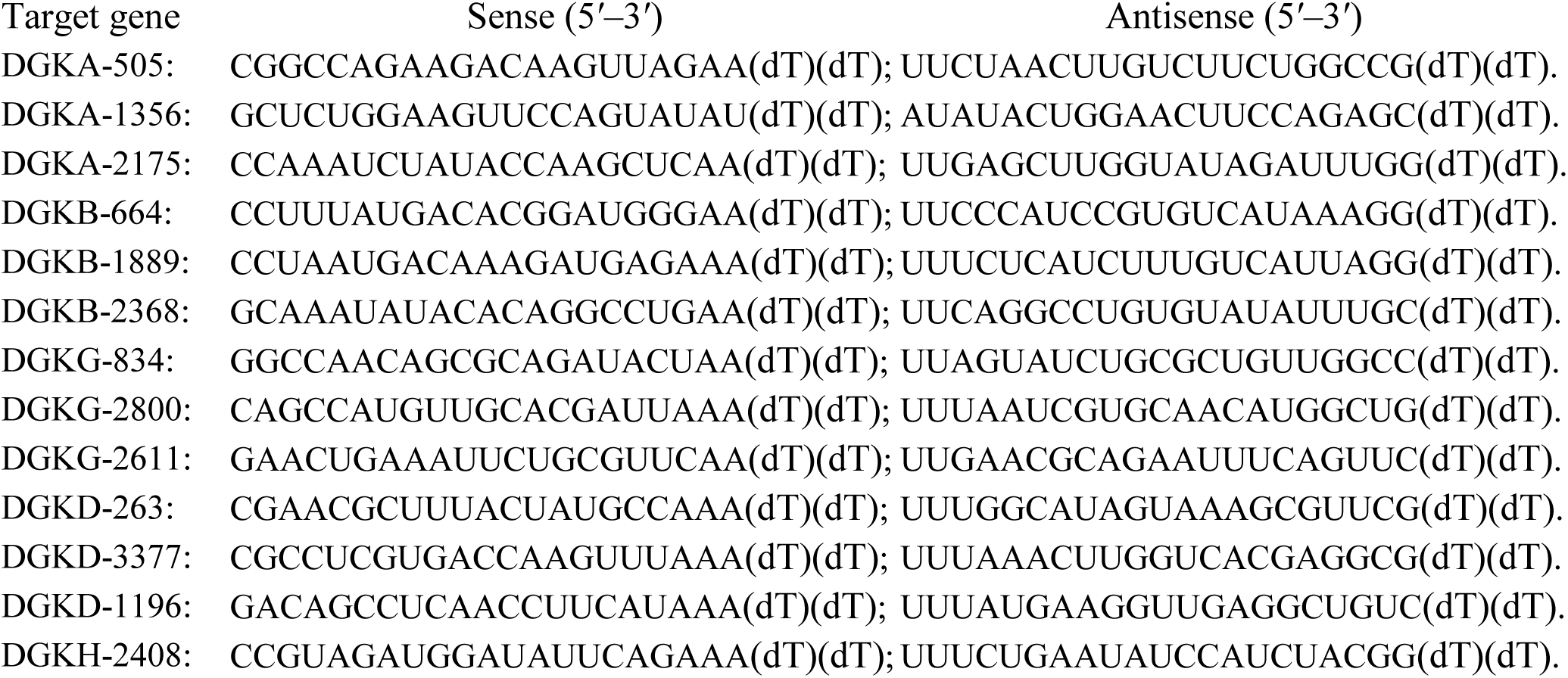
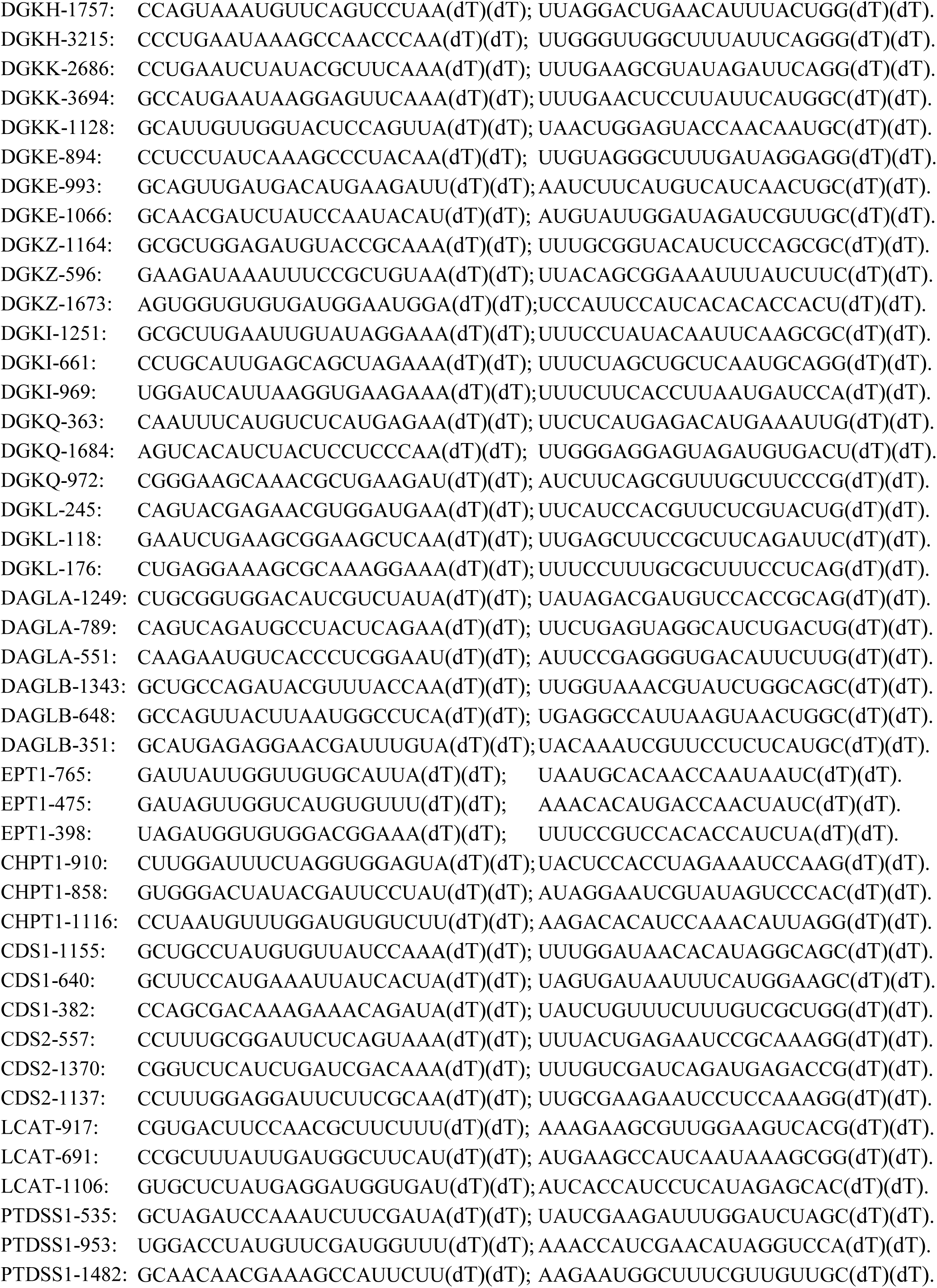
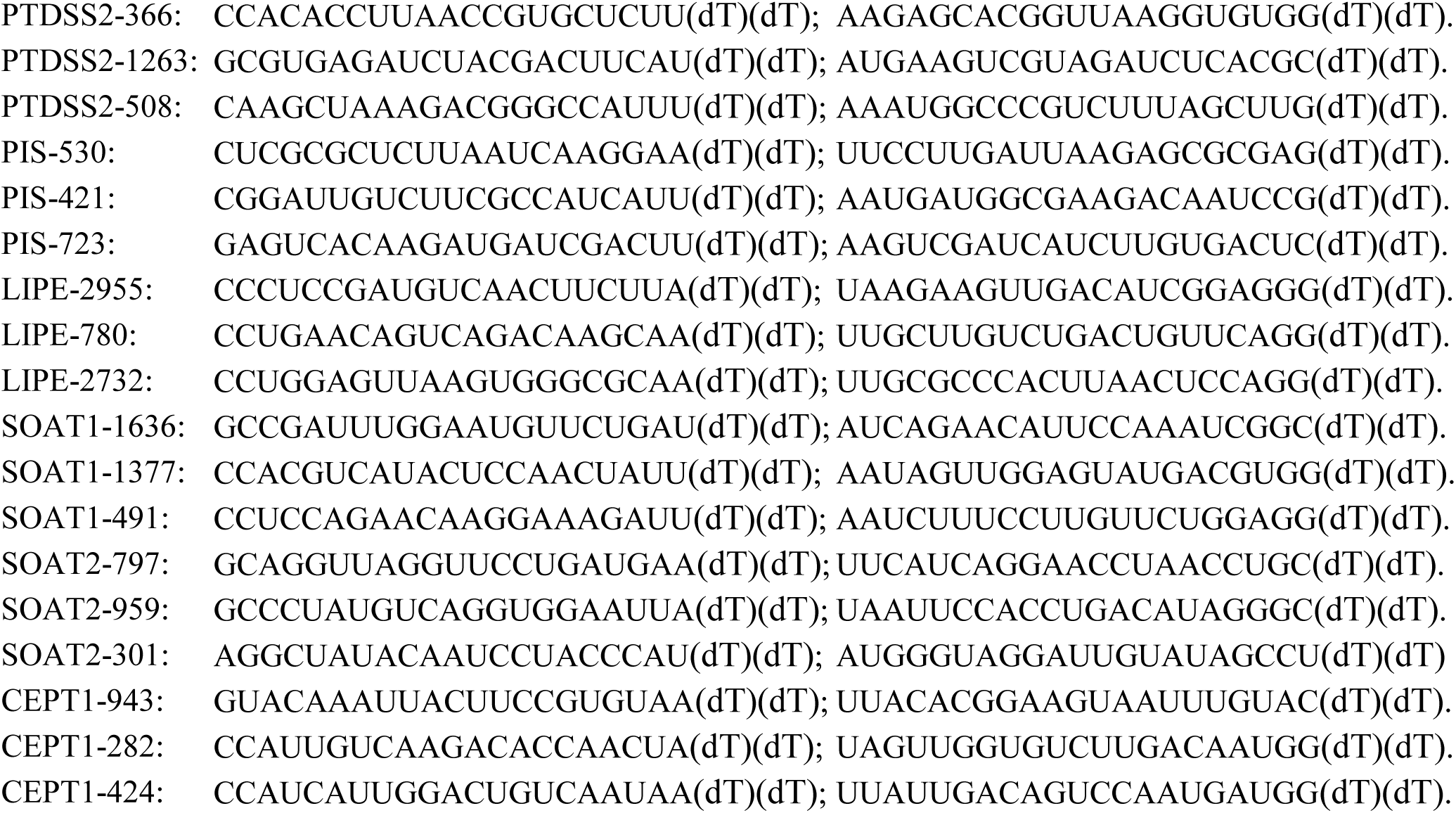

### qRT-PCR

qRT-PCR was performed following standard procedures. Briefly, total RNA was extracted using TRIzol reagent (Invitrogen, 15596026) and reverse-transcribed into cDNA with SuperScript II reverse transcriptase (Invitrogen, 18064014). qRT-PCR was carried out using SYBR GreenER qPCR SuperMix Universal (Invitrogen, 11762500), and reactions were run in technical triplicate on a Stratagene MX3000P qPCR system according to the manufacturer’s instructions. Threshold cycle (Ct) values for each gene were normalized to GAPDH, and relative expression levels were calculated using the 2^−ΔΔCt method.

The primers used in this study are listed below (5′–3′).

GAPDH: Forward, GGAGCGAGATCCCTCCAAAAT; Reverse, GGCTGTTGTCATACTTCTCATGG.
SOAT1: Forward, GAAGTTGGCAGTCACTTTGATGA; Reverse, GAGCGCACCCACCATTATCTA.
SOAT2: Forward, ATGGAAACACTGAGACGCACA; Reverse, GGTAGGATTGTATAGCCTCCCG.
(LIPE)HSL: Forward, TCAGTGTCTAGGTCAGACTGG; Reverse, AGGCTTCTGTTGGGTATTGGA.
DAGLA: Forward, CGGCCTGGTCTATAACCCG; Reverse, ATCTCAGCGATCATGCAGCTC.
DAGLB: Forward, ATGCCGGGGATGGTACTCTT; Reverse, CAGAATGCCAATCCACCACAG.
(CDIPT)PIS: Forward, GTGCCCAACCTCATCGGTTAT; Reverse, TGTCCATCGAAAGCGTCCAG.
LCAT: Forward, ACCTGGTCAACAATGGCTACG; Reverse, TAGAGCAAGTGTAGACAGCCG.
CDS1: Forward, AGTTCCTCATTCGCTACCATAGA; Reverse, GGTGTGACTGAGTGACAGTTATC.
CDS2: Forward, ATGGAGAGACTGCATCGGACA; Reverse, GAGGACCTCCGGGGTATCATC.
DGKK: Forward, GAGGCGACCTCAGAATCAGC; Reverse, GCTCTGGGACCGACTCTAGG.
DGKQ: Forward, TGAGCCAGACGCGGTTCTA; Reverse, CAGGCAACGTCCAACACTAC.
CEPT1: Forward, AACTAAAGCGGCTAGAAGAACAC; Reverse, CCATTCCCAATACCCTTGCAT.
CHPT1: Forward, CACCGAAGAGGCACCATACTG; Reverse, CCCTAAAGGGGAACAAGAGTTTG.
DGKD: Forward, CTTCGAGGGCGAACGCTTTA; Reverse, TTTTGGTACTGGATTCAGCTACG.
DGKE: Forward, GACGGGCACCTGATCTTGTG; Reverse, CTGGAGGCTACACCAGAAGG.
DGKZ: Forward, CGGAGGCCCCAGAATACTCT; Reverse, TTGTCGGGGATTGAGATACCA.
DGKB: Forward, CCTATCAGGCGGTCTGAGAAT; Reverse, GGAACACGTATTTGCAGGAGAAG.
DGKH: Forward, GATGCACAACTGGTACGCCT; Reverse, GGCCAGGGTAGTCCATTTACA.
PTDSS1: Forward, GCAAGTGGAGGACATCACCAT; Reverse, TCATCCCTGGTAAAGGCGAAG.
PTDSS2: Forward, CTCACCTGTACGCTTGGCTAT; Reverse, CCACAATACCTCTCTTGGTGTTG.

### LCAT activity assay

LCAT activity was measured using a fluorescence-based sterol esterification assay^30,31^. Briefly, phospholipid DMPC and dehydroergosterol (DHE) were mixed at a molar ratio of 9:1 to form a lipid film, which was subsequently rehydrated and sonicated in PBS buffer (pH 7.4) containing 1 mM EDTA. ApoA-I mimetic peptide or vehicle control was then added at a lipid-to-peptide mass ratio of 2:1, and synthetic HDL-like nanodiscs (sHDL-DHE) were generated through repeated thermal cycling (50 °C and 4 °C for 3 min each, repeated three times). For enzymatic reactions, sHDL-DHE nanodiscs containing varying concentrations of DHE (0–50 µM) were incubated with recombinant human LCAT (10 µg ml⁻¹); intracellular or medium-enriched LCAT was normalized to an equivalent concentration (10 µg ml⁻¹) based on immunoblot densitometry (Extended Data Fig. 9i). Reactions were carried out in PBS buffer supplemented with 1 mM EDTA and 60 µM bovine serum albumin at 37 °C with shaking for 30 min. The reactions were terminated by addition of stop solution containing cholesterol oxidase and Triton X-100, followed by further incubation at 37 °C for 1 h to selectively quench the fluorescence of unesterified DHE. Fluorescence corresponding to esterified DHE was measured using a microplate reader at an excitation wavelength of 325 nm and an emission wavelength of 425 nm. LCAT activity was calculated based on a DHE standard curve and expressed as the amount of DHE ester generated per unit enzyme per unit time.

### Immunofluorescence microscopy

HT-1080 and SK-HEP-1 cells were seeded onto glass-bottom 35 mm dishes and, where indicated, transfected with CHPT1-GFP or CHPT1-3×Flag plasmids. Upon reaching 70–80% confluence, cells were washed twice with PBS and fixed with 4% paraformaldehyde for 10 min at room temperature, followed by two additional PBS washes. Cells were permeabilized with either 0.1% Triton X-100 or 10 µg ml⁻¹ digitonin for 10 min and subsequently blocked with a protein-free rapid blocking solution (Yeasen, PS108P) for 15 min.

Cells were then incubated overnight at 4 °C with the indicated primary antibodies: anti-LCAT (Proteintech, cat. no. 12243-1-AP; 1:200), anti-TGN46 (Proteintech, cat. no. 83099-2-RR; 1:200), anti-GM130 (Abcam, cat. no. ab52649; 1:400), anti-golgin 97 (Proteintech, cat. no. 68648-1-Ig; 1:200), anti-Flag (Proteintech, cat. no. 20543-1-AP; 1:500), and Alexa Fluor 647-conjugated anti-GRP94 (Abcam, cat. no. ab52649; 1:1,000). After washing, cells were incubated for 1 h at room temperature in the dark with fluorophore-conjugated secondary antibodies: anti-rabbit (H+L)/TRITC (ZSGB-BIO, cat. no. ZF-0316; 1:200) and anti-mouse (H+L)/FITC (ZSGB-BIO, cat. no. ZF-0311; 1:200). Nuclear DNA was counterstained with DAPI.

Images were acquired using a Zeiss LSM 880 confocal microscope equipped with a 63× oil-immersion objective and analysed using ZEN 2.3 SP1 software.

### Lipid peroxidation assay

Cells were treated with the indicated compounds for the specified times, collected by trypsinization, and resuspended in 200 µl PBS containing 5 µM C11-BODIPY 581/591 (Invitrogen). Cells were incubated at 37 °C for 30 min in a water bath and subsequently analysed on a BD Accuri C6 flow cytometer using a 488 nm excitation laser, with lipid peroxidation detected in the FL1 channel. At least 10,000 single cells were analysed per sample.

### Tumour xenografts in nude mice

Female athymic nude mice (4–5 weeks old) were purchased from Beijing Vita River Laboratory Animal Technology Co. Mice were housed in a pathogen-free animal facility according to the principles and procedures outlined in the NIH Guide for the Care and Use of Laboratory Animals. The animals were maintained under the following conditions: temperature: 19–23°C; humidity: 40–70%; light/dark cycle: 12/12 hours. The maximum tumour burden allowed by the ethical committee was less than 10% of the body weight, which was not exceeded in this study. A total of 2 × 10⁶ HT-1080 cells or 8 × 10⁶ SK-HEP-1 cells were subcutaneously injected into the mice. Once the tumours reached 100–200 mm³, mice were randomly assigned to different treatment groups. Drugs were dissolved in a solution of 10% dimethyl sulfoxide (DMSO), 40% polyethylene glycol (PEG) 300, 5% Tween 80, and 45% saline, and administered via intraperitoneal injection at the following doses: SAS: 100 mg kg^-1^, JW642: 20 mg kg^-1^, Lip-1: 10 mg kg^-1^, DO34: 20 mg kg^-1^, and KT109: 20 mg kg^-1^. Treatment was administered daily. Tumour volume was calculated using the formula:Volume (mm³) = Length (mm) × (Width (mm))² × 1/2.

### 4-HNE Immunofluorescence Staining

Excised tumours were fixed in 4% paraformaldehyde (PFA) for 24 h at room temperature. Following fixation, tissue samples were dehydrated in a graded ethanol series, cleared with xylene, and embedded in paraffin. Sections (3–5 μm) were prepared for immunofluorescence staining. Tumour sections were first blocked with 3% hydrogen peroxide (H₂O₂) for 30 min at room temperature to quench endogenous peroxidase activity. Antigen retrieval was performed using 1 M sodium citrate buffer (pH 6.5) for 30 min at 95°C. After washing with PBS, sections were incubated overnight at 4°C with anti-4-HNE antibody (1:400, Abcam, cat. no. ab46545). The following day, sections were washed and incubated with HRP-conjugated anti-rabbit IgG (1:200, Beijing Biodragon Immunotechnologies, cat. no. BF03008) for 1 h at room temperature. Staining was visualized using DAB solution and counterstained with hematoxylin. Images were acquired and analysed to quantify the percentage of positive staining.

### MDA quantification in liver tissue and cultured cells

Liver tissues or cells were homogenized or lysed in PBS or Western/IP lysis buffer (P0013, Beyotime), with tissue samples prepared at a ratio of 10% (w/v) and cell samples processed with 0.1 ml lysis buffer per 1 × 10⁶ cells. Homogenates were centrifuged at 10,000–12,000 g for 10 min to collect the supernatant, and samples were further clarified through a 0.2 µm filter when necessary. All procedures were performed on ice or at 4 °C. Protein concentrations were determined using the BCA Protein Assay Kit. For MDA measurement, 0.1 ml of the sample or standard solution was mixed with 0.2 ml of MDA working reagent, heated at 100 °C for 15 min, cooled to room temperature, and centrifuged at 1,000 g for 10 min. A total of 200 µl of the resulting supernatant was transferred to a 96-well plate, and absorbance was measured at 532 nm. MDA levels were calculated using a standard curve and normalized to protein content or tissue weight, expressed as μmol mg⁻¹ protein or μmol mg⁻¹ tissue.

### MASH mouse model and endpoint analysis

Six-week-old C57BL/6J mice were purchased from Vital River and fed an AMLN diet to induce MASH. After 12 weeks of feeding, mice received tail-vein injections of GalNAc-siRNA (2 mg kg⁻¹) every three days while continuing on the AMLN diet. Body weight, food intake and water intake were recorded every three days throughout the study. At week 26, mice were euthanized for endpoint analyses. Serum was collected for biochemical measurements, including ALT, AST, TC, TAG, HDL-C and LDL-C. Liver tissue and adipose depots were dissected, weighed and photographed. Liver samples were used to measure TAG, TC, MDA and cholesteryl ester levels, and siRNA knockdown efficiency was assessed by Western blotting. Portions of the liver were processed for paraffin or frozen sections and subjected to H&E, Oil Red O and Sirius Red staining, as well as 4-HNE and CD45 immunofluorescence, to evaluate steatosis, inflammation, fibrosis and ferroptosis levels.

The GalNAc-siRNA sequences used for *in vivo* knockdown were as follows (sense and antisense strands, 5′–3′). CHPT1#1: Sense, CCUGGACUCCACAUAGGAUUA (dT)(dT); Antisense, UAAUCCUAUGUGGAGUCCAGG(dT)(dT). CHPT1#2: Sense, GCACCAUACUGGACAUACCUU(dT)(dT); Antisense, AAGGUAUGUCCAGUAUGGUGC(dT)(dT). LCAT#1: Sense, GCAAUAAGACACUGGAGCA(dT)(dT); Antisense, UGCUCCAGUGUCUUAUUGC(dT)(dT). LCAT#2: Sense, ACAUAAAG CUGAAAGAGGA(dT)(dT); Antisense, UCCUCUUUCAGCUUUAUGU(dT)(dT). A non-targeting siRNA was used as a negative control (siControl): Sense, UUCUCCGA CAGUGUCACGU(dT)(dT); Antisense, ACGUGACACUGUCGGAGAA(dT)(dT).

### Statistics and reproducibility

All experiments were independently repeated as indicated, with *n* denoting the number of biological replicates. Samples were randomly assigned to experimental groups, and investigators were blinded to group allocation during data collection and analysis. No data were excluded from the analyses. Sample sizes were not predetermined by statistical methods but were chosen to be comparable to those used in previous studies^48,49^. Data distribution was assumed to be normal but was not formally tested. Results are presented as mean ± s.d. Statistical analyses and graphing were performed using Prism 7 (GraphPad Software). Statistical significance was determined using one-way analysis of variance (ANOVA), as specified in the figure legends.

## Acknowledgements

We thank Dr. Xiaohui Liu (Metabolomics Facility at Tsinghua University Branch of China National Center for Protein Sciences, China) for technical help. This work was supported by grants from the National Natural Science Foundation of China (82325038 and 82530094), and the Chinese Institutes for Medical Research, Beijing (CX23YZ18).

## Author contributions

B.L. conceived the study and designed the experiments. L.M. and P.T. performed the experiments. Q.Z., Q.L., J.L., H.W. and Y.Z. generated plasmid constructs and established cell lines. L.M., Z.G. and R.Y. collected and analysed the data. L.Q., Y.F. and L.L. provided conceptual advice and technical support. L.M. developed and optimized the experimental methodologies. L.M. and B.L. wrote the manuscript, and L.M., P.T. and Q.Z. edited the manuscript.

## Extended Data Figure Legends

**Extended Data Fig. 1 | MAGL inhibitors promote ferroptosis. a**, Schematic illustration of the viability-based chemical screen using a small-molecule library targeting metabolic enzymes; compounds were applied at 10 µM to HT-1080 cells pretreated with DMSO or ferrostatin-1 (Fer-1, 5 µM) for 12 h, followed by cell viability assessment after 72 h. **b**, Cell viability of HT-1080 cells treated with increasing concentrations of indicated MAGL inhibitors (JW642, JZL184, Elcubragis (Elcu), JJKK048 (JK048) or MAGL-IN-1 (MI-1)) for 24 h in the presence of DMSO or Fer-1 (5 µM). **c**, Malondialdehyde (MDA) levels in HT-1080 cells treated with or without JW642 or JZL184 (20 µM) for 12 h in the presence or absence of Fer-1 (5 µM). **d**, Lipid peroxidation levels in HT-1080 cells treated with DMSO, RSL3 (10 nM, positive control), JW642 or JZL184 (20 µM) for 24 h in the presence or absence of Fer-1 (5 µM). **e**, Representative transmission electron microscopy images showing mitochondrial morphology in HT-1080 cells treated with DMSO, JW642 or JZL184 (20 µM) for 24 h in the presence or absence of Fer-1 (5 µM). Scale bar, 2 µm. **f**,**g**, Cell viability of HT-1080 cells treated with increasing concentrations of RSL3 (**f**) or erastin (**g**) in the presence of DMSO or 10 µM MAGL inhibitors (JW642, JZL184, Elcu, JK048 or MI-1) for 24 h. **h**,**i**, Cell viability of HT-1080 cells treated with increasing concentrations of RSL3 or erastin in the presence of DMSO, JW642 or JZL184 (10 µM), in combination with Fer-1 (5 µM), liproxstatin-1 (Lip-1, 10 µM), Z-VAD-FMK (Z-VAD, 10 µM) or necrostatin-1 (Nec-1, 10 µM). Data in **b–d** and **f–i** are mean ±s.d. from three biological replicates (n = 3). Statistical analysis in **c** and **d** was performed using one-way ANOVA; *P < 0.05, **P < 0.01, ***P < 0.001.

**Extended Data Fig. 2 | ATGL inhibition attenuates ferroptosis. a**,**b**, Immunoblot analysis confirming efficient knockout of MAGL or ATGL in HT-1080 (**a**) and NCI-H226 (**b**) cells using two independent sgRNAs for each gene. **c**, Cell viability of NCI-H226 cells with or without MAGL or ATGL knockout following treatment with increasing concentrations of RSL3 or erastin for 24 h. **d,e**, Lipid peroxidation levels in HT-1080 (**d**) and NCI-H226 (**e**) cells with or without MAGL knockout following treatment with DMSO, RSL3 or erastin for 4 h; RSL3 was used at 10 nM and erastin at 5 µM in HT-1080 cells (**d**), whereas RSL3 was used at 1 µM and erastin at 20 µM in NCI-H226 cells (**e**). **f**, Lipid peroxidation levels in NCI-H226 cells with or without ATGL knockout following treatment with DMSO, RSL3 (5 µM) or erastin (40 µM) for 4 h. **g**,**h**, Cell viability of HT-1080 (**g**) and NCI-H226 (**h**) cells treated with increasing concentrations of RSL3 or erastin in the presence of DMSO, ATGL inhibitor atglistatin (10 µM) or MAGL inhibitors (JW642 or JZL184 at 10 µM) for 24 h. Data in **c–h** are mean ±s.d. from three biological replicates (n = 3). Statistical analysis in **d–f** was performed using one-way ANOVA; ns, not significant; *P < 0.05, **P < 0.01, ***P < 0.001.

**Extended Data Fig. 3 | DAGL inhibitors sensitize cancer cells to ferroptosis. a**, Cell viability of HT-1080 and NCI-H226 cells treated with increasing concentrations of RSL3 or erastin in the presence of DMSO or the hormone-sensitive lipase (HSL) inhibitors HSL-IN-1 or HSL-IN-3 (10 µM). **b**, Lipid peroxidation levels and quantitative PCR (qPCR) analysis of HSL expression in HT-1080 and NCI-H226 cells transfected with control or HSL-targeting siRNA. **c**, Cell viability of a panel of cancer cell lines (HT-1080, SK-HEP-1, HeLa, NCI-H226, MCF-7, A549, HepG2, SNU878 and HCT116) treated with increasing concentrations of RSL3 in the presence of DMSO or the DAGL inhibitor DO34 (10 µM). **d–i**, Cell viability of HT-1080 (**d–f**) and SK-HEP-1 (**g–i**) cells treated with increasing concentrations of RSL3 (**d**,**g**) or erastin (**e**,**f**,**h**,**i**) in the presence of DMSO or the DAGL inhibitors KT109 (10 µM) or DO34 (10 µM). Data in **a–i** are mean ±s.d. from three biological replicates (n = 3).

**Extended Data Fig. 4 | DAG determines the sensitization of cancer cells to ferroptosis. a**,**b**, Immunoblot analysis confirming efficient knockout of DAGLA or DAGLB in HT-1080 (**a**) and SK-HEP-1 (**b**) cells using two independent sgRNAs for each gene. **c**, Cell viability of SK-HEP-1 cells with or without DAGLB knockout following treatment with increasing concentrations of RSL3 or erastin for 24 h. **d**, Lipid peroxidation levels in SK-HEP-1 cells with or without DAGLB knockout following treatment with DMSO, RSL3 (10 nM) or erastin (5 µM) for 4 h. **e**, Cell viability of HT-1080 cells overexpressing empty vector, DAGLA or DAGLB following treatment with increasing concentrations of RSL3 or erastin for 24 h. Immunoblot analysis confirming overexpression of DAGLA or DAGLB (top). **f**, Lipid peroxidation levels in HT-1080 cells overexpressing vector, DAGLA or DAGLB following treatment with DMSO, RSL3 (20 nM) or erastin (10 µM) for 4 h. **g**, Cell viability of SK-HEP-1 cells overexpressing vector, DAGLA or DAGLB following treatment with increasing concentrations of RSL3 or erastin for 24 h. Immunoblot analysis confirming overexpression of DAGLA or DAGLB (top). **h**, Lipid peroxidation levels in SK-HEP-1 cells overexpressing vector, DAGLA or DAGLB following treatment with DMSO, RSL3 (20 nM) or erastin (10 µM) for 4 h. **i**,**j**, Cell viability of HT-1080 (**i**) and SK-HEP-1 (**j**) cells treated with increasing concentrations of DO34 or KT109 for 24 h in the presence of DMSO or atglistatin (10 µM). **k**, Immunoblot analysis confirming efficient knockout of ATGL in SK-HEP-1 cells. **l**, Cell viability of SK-HEP-1 cells with or without ATGL knockout treated with increasing concentrations of DO34 or KT109 for 24 h. **m**,**n**, Lipid peroxidation levels in HT-1080 (**m**) and SK-HEP-1 (**n**) cells with or without ATGL knockout after treatment with DMSO, DO34 (20 µM) or KT109 (20 µM) for 4 h. Data in **c–j** and **l–n** are mean ±s.d. from three biological replicates (n = 3). Statistical analysis in **d**, **f**, **h**, **m** and **n** was performed using one-way ANOVA; ns, not significant; *P < 0.05, **P < 0.01, ***P < 0.001.

**Extended Data Fig. 5 | DAGL inhibition induces ferroptosis. a**,**b**, Cell viability of a panel of cancer cell lines treated with increasing concentrations of the DAGL inhibitors DO34 (**a**) or KT109 (**b**) for 24 h in the presence of DMSO or ferrostatin-1 (Fer-1, 5 µM). **c**, Cell viability of SK-HEP-1 cells treated with increasing concentrations of DO34 or KT109 for 24 h in the presence of DMSO, Fer-1 (5 µM), necrostatin-1 (Nec-1, 10 µM) or Z-VAD-FMK (Z-VAD, 10 µM). **d**, MDA levels in HT-1080 cells treated with or without DO34 or KT109 (20 µM) for 12 h in the presence or absence of Fer-1 (5 µM). e,**f**, Lipid peroxidation levels in HT-1080 (**e**) and SK-HEP-1 (**f**) cells treated with DMSO, DO34 or KT109 (20 µM) for 4 h in the presence or absence of Fer-1 (5 µM). Data in **a–f** are mean ±s.d. from three biological replicates (n = 3). Statistical analysis in d–f was performed using one-way ANOVA; ns, not significant; *P < 0.05, **P < 0.01, ***P < 0.001.

**Extended Data Fig. 6 | Cholesteryl ester accumulation is a causal determinant of ferroptosis. a**, Cell viability of HT-1080 cells treated with increasing concentrations of DO34 or KT109 in the presence of DMSO or the PKC inhibitor GF109203X (10 µM) for 24 h. **b**, Lipid peroxidation levels in HT-1080 cells treated with DO34 or KT109 in the presence of DMSO or GF109203X (10 µM) for 4 h. **c**, Lipidomics profiling of selected ChE species in HT-1080 cells treated with DMSO, atglistatin (10 µM) or DO34 (20 µM) for 12 h. **d**, Lipidomics profiling of selected ChE species in HT-1080 cells treated with DMSO, atglistatin (10 µM) or JW642 (20 µM) for 12 h. **e**, Cell viability of SK-HEP-1 cells treated with increasing concentrations of ChE(20:4) or ChE(18:2) for 24 h in the presence of DMSO or Fer-1 (5 µM). **f**,**g**, Cell viability of SK-HEP-1 and HT-1080 cells treated with increasing concentrations of RSL3 or erastin in the presence of DMSO or exogenous ChE(20:4) or ChE(18:2) (20 µM) for 24 h. **h**, Lipid peroxidation levels in SK-HEP-1 cells treated with or without ChE(18:2) or ChE(20:4) (20 µM) for 4 h in the presence or absence of Fer-1 (5 µM). **i**,**j**, MDA levels in HT-1080 (**i**) and SK-HEP-1 (**j**) cells treated with or without ChE(18:2) or ChE(20:4) (20 µM) for 12 h in the presence or absence of Fer-1 (5 µM). **k**,**l**, Cell viability of HT-1080 and SK-HEP-1 cells with or without ATGL knockout treated with increasing concentrations of RSL3 or erastin for 24 h in the presence or absence of ChE(20:4) (**k**) or ChE(18:2) (**l**) (20 µM). Data in **b–l** are mean ±s.d. from three biological replicates (n = 3). Statistical analysis in **h–j** was performed using one-way ANOVA; ns, not significant; *P < 0.05, **P < 0.01, ***P < 0.001.

**Extended Data Fig. 7 | siRNA screening identifies CHPT1 and LCAT as mediators of DAGL inhibition-induced ferroptosis. a**, Lipid peroxidation levels in HT-1080 and SK-HEP-1 cells transfected with siRNAs targeting SOAT1 or SOAT2, or siControl, following treatment with DMSO or DO34 (20 µM) for 4 h. **b**,**c**, Quantitative RT-PCR analysis confirming knockdown efficiency of SOAT1 and SOAT2 in HT-1080 (**b**) and SK-HEP-1 (**c**) cells. **d**, Heat map showing relative mRNA expression levels (Z-score normalized) of genes involved in phospholipid and DAG metabolism following siRNA transfection in HT-1080 cells. **e–m**, Lipid peroxidation-based functional screening in HT-1080 cells following siRNA-mediated knockdown of DAG-to-phospholipid conversion enzymes, including DAGLA, DAGLB, DGK family members, CDS1/2, PTDSS1/2, CEPT1, EPT1, PIS, LCAT and CHPT1, with cells treated with DMSO or DO34 (20 µM) for 4 h. Data in **a–m** are mean ±s.d. from three biological replicates (n = 3). Statistical analysis in **a** and **e–m** was performed using one-way ANOVA; ns, not significant; *P < 0.05, **P < 0.01, ***P < 0.001.

**Extended Data Fig. 8 | The CHPT1–LCAT axis regulates DAG routing to ferroptosis. a**,**b**, Immunoblot analysis confirming efficient knockout of CHPT1 or LCAT in HT-1080 (**a**) and SK-HEP-1 (**b**) cells using two independent sgRNAs for each gene. **c**,**d**, Lipid peroxidation levels in HT-1080 and SK-HEP-1 cells with or without knockout of LCAT (**c**) or CHPT1 (**d**) following treatment with DMSO, RSL3 (20 nM) or erastin (10 µM) for 4 h. **e**,**f**, Lipid peroxidation levels in SK-HEP-1 cells with or without knockout of LCAT (**e**) or CHPT1 (**f**) following treatment with DMSO, DO34 or KT109 (20 µM) for 4 h. **g**, Cell viability of SK-HEP-1 cells with or without knockout of LCAT or CHPT1 following treatment with increasing concentrations of DO34 or KT109 for 24 h. **h**,**i**, Cell viability of HT-1080 and SK-HEP-1 cells with or without knockout of LCAT or CHPT1 following treatment with increasing concentrations of RSL3 (**h**) or erastin (**i**) for 24 h. Data in **c–i** are mean ± s.d. from three biological replicates (n = 3). Statistical analysis in **c–f** was performed using one-way ANOVA; ns, not significant; *P < 0.05, **P < 0.01, ***P < 0.001.

**Extended Data Fig. 9 | Intracellular LCAT mediates ferroptosis. a**,**b**, Immunoblot analysis confirming expression of wild-type LCAT–Flag, signal peptide-deficient ΔLCAT–Flag or catalytically inactive LCAT-S205A–Flag in HT-1080 (**a**) and SK-HEP-1 (**b**) cells. **c**, Immunoblot analysis of LCAT in cell lysates and Flag-enriched conditioned medium from HT-1080 cells expressing the indicated constructs; recombinant LCAT (reLCAT) purified from *Escherichia coli* was included as a control. **d–f**, Cell viability of SK-HEP-1 (**d**,**f**) and HT-1080 (**e**) cells expressing the indicated constructs following treatment with increasing concentrations of DO34 or KT109 (**d**), or RSL3 or erastin (**e**,**f**) for 24 h. **g**,**h**, Lipid peroxidation levels in HT-1080 (**g**) and SK-HEP-1 (**h**) cells expressing the indicated constructs following treatment with DMSO, RSL3 (10 nM), erastin (5 µM), DO34 (10 µM) or KT109 (10 µM) for 4 h. **i**, Immunoblot analysis following Flag-bead purification of LCAT from cell lysates and conditioned medium of transfected HEK293T cells. reLCAT (recombinant LCAT) purified from *Escherichia coli* was included as a control. **j**, *In vitro* DHE acyltransferase activity of purified LCAT–Flag proteins. Activity of wild-type, ΔLCAT and catalytically inactive S205A LCAT–Flag proteins was measured across increasing concentrations of DHE in the presence or absence of ApoA-I. Recombinant LCAT (reLCAT), LCAT–Flag recovered from cell lysates (C) and LCAT–Flag enriched from conditioned medium (secreted fraction; M) were analysed. Detailed procedures are described in Methods. Data obtained in the absence of ApoA-I are also summarized in Fig. 3c. Data in **d–h** and j are mean ±s.d. from three biological replicates (n = 3). Statistical analysis in **g** and **h** was performed using one-way ANOVA; ns, not significant; *P < 0.05, **P < 0.01, ***P < 0.001.

**Extended Data Fig. 10 | Intracellular localization of CHPT1 and LCAT. a**, Immunofluorescence images showing localization of CHPT1–Flag with Golgi markers (GM130, Golgin-97 and TGN46) and GRP78 (an endoplasmic reticulum luminal protein marker) following selective permeabilization with 0.1% Triton X-100 or 10 µg ml⁻¹digitonin. Scale bar, 2 µm. **b**, Immunofluorescence images showing co-localization of LCAT, CHPT1–EGFP, GRP78 and TGN46 following selective permeabilization with 0.1% Triton X-100 or 10 µg ml⁻¹digitonin. Scale bar, 2 µm.

**Extended Data Fig. 11 | CHPT1–LCAT mediates ferroptosis across diverse contexts. a**, Immunoblot analysis of CHPT1 and LCAT protein expression across the indicated human cancer cell lines. **b**, Heat maps showing relative mRNA levels (Z-score-normalized) of CHPT1 (left) or LCAT (right) following siRNA-mediated knockdown across the indicated cell lines. **c**, Lipid peroxidation levels in the indicated cell lines transfected with control or LCAT-targeting siRNA and treated with DMSO, DO34 or KT109 (20 µM). **d**, Lipid peroxidation levels in the indicated cell lines transfected with control or CHPT1-targeting siRNA and treated with DMSO, DO34 or KT109 (20 µM). Data in **c** and **d** are mean ±s.d. from three biological replicates (n = 3). Statistical analysis in **c** and **d** was performed using one-way ANOVA; ns, not significant; *P < 0.05, **P < 0.01, ***P < 0.001.

**Extended Data Fig. 12 | MAGL and DAGL inhibitors suppress tumour growth via ferroptosis. a**, Representative images of excised tumours from HT-1080 xenografts treated with DMSO or the MAGL inhibitor JW642 alone or in combination with SAS and/or Lip-1 at the experimental endpoint. **b**, Tumour growth curves of HT-1080 xenografts measured over time under the indicated treatment conditions shown in **a**. **c**, Representative images of excised tumours from HT-1080 xenografts treated with DMSO or DAGL inhibitors (DO34 or KT109) alone or in combination with SAS and/or Lip-1 at the experimental endpoint. **d**, Tumour growth curves of HT-1080 xenografts measured over time under the indicated treatment conditions shown in **c**. **e**, Representative images of excised tumours from SK-HEP-1 xenografts treated with DMSO or DAGL inhibitors (DO34 or KT109) alone or in combination with SAS and/or Lip-1 at the experimental endpoint. **f**, Tumour growth curves of SK-HEP-1 xenografts measured over time under the indicated treatment conditions shown in **e**. Data in **b**, **d** and **f** are mean ±s.d. n = 6 (**a**,**b**), 8 (**c**,**d**) or 7 (**e**,**f**) tumours. All xenograft experiments were performed in immunodeficient mice.

**Extended Data Fig. 13 | GalNAc–siRNA enables hepatic knockdown of CHPT1 and LCAT in mice. a**, Schematic illustration of GalNAc–siRNA-mediated hepatocyte targeting via the asialoglycoprotein receptor (ASGPR), showing receptor-mediated endocytosis, endosomal release of siRNA, loading into the RNA-induced silencing complex (RISC) and ASGPR recycling. **b**, Immunoblot analysis of liver extracts from mice treated with GalNAc-conjugated siRNAs targeting CHPT1 or LCAT, or a GalNAc–siControl, demonstrating efficient hepatic knockdown of CHPT1 (left) and LCAT (right); Lamin B and GAPDH serve as loading controls, respectively.

**Extended Data Fig. 14 | Targeting the CHPT1–LCAT axis attenuates MASH in mice. a–d**, Serum triacylglycerols (TAG) (**a**), total cholesterol (TC) (**b**), HDL cholesterol (HDL-C) (**c**) and LDL cholesterol (LDL-C) (**d**) measured at the experimental endpoint in normal-diet-fed (ND) and MASH mice treated with GalNAc–siControl or GalNAc–siRNAs targeting CHPT1 or LCAT. **e**,**f**, Hepatic TAG (**e**) and TC (**f**) contents. **g**,**h**, Liver weight (**g**) and liver-to-body-weight ratio (**h**). **i–l**, Weights of adipose depots, including retroperitoneal white adipose tissue (rWAT) (**i**), epididymal white adipose tissue (eWAT) (**j**), inguinal white adipose tissue (iWAT) (**k**) and brown adipose tissue (BAT) (**l**). **m**,**n**, Representative Oil Red O staining of liver sections (**m**) and corresponding quantification of Oil Red O-positive area (**n**). Scale bar, 100 µm. Quantification in **n** was performed from eight mice per group with three randomly selected fields per mouse. **o**, Food intake, water intake and body-weight trajectories over the course of the experiment. Data in **a–l** and **n** are mean ±s.d. from eight mice per group. Statistical analysis in a–l and n was performed using one-way ANOVA; ns, not significant; *P < 0.05, **P < 0.01, ***P < 0.001.

**Extended Data Fig. 15 | DAG routing drives ferroptosis via a CHPT1–LCAT metabolic axis.** Lipolytic hydrolysis of triacylglycerol (TAG) by ATGL generates diacylglycerol (DAG), which is normally further metabolized by DAGL and MAGL. When DAG availability is increased, a distinct intracellular lipid-routing pathway is engaged, whereby DAG is selectively channelled into a metabolic axis composed of CHPT1 and an enzymatically active intracellular pool of LCAT (iLCAT) localized to Golgi and trans-Golgi network (TGN) membranes. This axis converts DAG into phosphatidylcholine (PC) and polyunsaturated cholesteryl esters (PUFA-ChE), promoting lipid reactive oxygen species (Lipid ROS) accumulation and ferroptotic cell death. In parallel, the canonical serum-active LCAT (here termed eLCAT) traffics through the ER–Golgi secretory pathway to exert its extracellular functions. This schematic illustrates that ferroptotic vulnerability is dictated by subcellular lipid routing rather than bulk lipolytic flux and highlights the dual functional role of the CHPT1–LCAT axis in intracellular lipid metabolism and execution of ferroptosis.

## Reference

1 Dixon, S. J. et al. Ferroptosis: an iron-dependent form of nonapoptotic cell death. Cell 149, 1060–1072 (2012). 10.1016/j.cell.2012.03.042

2 Lei, G., Zhuang, L. & Gan, B. The roles of ferroptosis in cancer: Tumor suppression, tumor microenvironment, and therapeutic interventions. Cancer Cell 42, 513–534 (2024). 10.1016/j.ccell.2024.03.011

3 Peleman, C., Francque, S. & Berghe, T. V. Emerging role of ferroptosis in metabolic dysfunction-associated steatotic liver disease: revisiting hepatic lipid peroxidation. EBioMedicine 102, 105088 (2024). 10.1016/j.ebiom.2024.105088

4 Yang, C. et al. De novo pyrimidine biosynthetic complexes support cancer cell proliferation and ferroptosis defence. Nat Cell Biol 25, 836–847 (2023). 10.1038/s41556-023-01146-4

5 Dixon, S. J. & Olzmann, J. A. The cell biology of ferroptosis. Nat Rev Mol Cell Biol 25, 424–442 (2024). 10.1038/s41580-024-00703-5

6 Pope, L. E. & Dixon, S. J. Regulation of ferroptosis by lipid metabolism. Trends Cell Biol 33, 1077–1087 (2023). 10.1016/j.tcb.2023.05.003

7 Allevato, M. M. et al. A genome-wide CRISPR screen reveals that antagonism of glutamine metabolism sensitizes head and neck squamous cell carcinoma to ferroptotic cell death. Cancer Lett 598, 217089 (2024). 10.1016/j.canlet.2024.217089

8 Lee, H. et al. Energy-stress-mediated AMPK activation inhibits ferroptosis. Nat Cell Biol 22, 225–234 (2020). 10.1038/s41556-020-0461-8

9 Fan, F. et al. A Dual PI3K/HDAC Inhibitor Induces Immunogenic Ferroptosis to Potentiate Cancer Immune Checkpoint Therapy. Cancer Res 81, 6233–6245 (2021). 10.1158/0008-5472.CAN-21-1547

10 Nomura, D. K. et al. Monoacylglycerol lipase regulates a fatty acid network that promotes cancer pathogenesis. Cell 140, 49–61 (2010). 10.1016/j.cell.2009.11.027

11 Chang, J. W. et al. Highly selective inhibitors of monoacylglycerol lipase bearing a reactive group that is bioisosteric with endocannabinoid substrates. Chem Biol 19, 579–588 (2012). 10.1016/j.chembiol.2012.03.009

12 Xie, Y. et al. Ferroptosis: process and function. Cell Death Differ 23, 369–379 (2016). 10.1038/cdd.2015.158

13 Friedmann Angeli, J. P., et al. Inactivation of the ferroptosis regulator Gpx4 triggers acute renal failure in mice. Nat Cell Biol 16, 1180–1191 (2014). 10.1038/ncb3064

14 Zilka, O. et al. On the Mechanism of Cytoprotection by Ferrostatin-1 and Liproxstatin-1 and the Role of Lipid Peroxidation in Ferroptotic Cell Death. ACS Cent Sci 3, 232–243 (2017). 10.1021/acscentsci.7b00028

15 Slee, E. A. et al. Benzyloxycarbonyl-Val-Ala-Asp (OMe) fluoromethylketone (Z-VAD.FMK) inhibits apoptosis by blocking the processing of CPP32. Biochem J 315 **( Pt** 1), 21–24 (1996). 10.1042/bj3150021

16 Degterev, A. et al. Chemical inhibitor of nonapoptotic cell death with therapeutic potential for ischemic brain injury. Nat Chem Biol 1, 112–119 (2005). 10.1038/nchembio711

17 Zimmermann, R. et al. Fat mobilization in adipose tissue is promoted by adipose triglyceride lipase. Science 306, 1383–1386 (2004). 10.1126/science.1100747

18 Taschler, U. et al. Monoglyceride lipase deficiency in mice impairs lipolysis and attenuates diet-induced insulin resistance. J Biol Chem 286, 17467–17477 (2011). 10.1074/jbc.M110.215434

19 Motamedi, S. et al. AMP-activated protein kinase-driven lipid droplet dynamics govern melanoma sensitivity to polyunsaturated fatty acid and iron-induced ferroptosis. Nat Commun 16, 11252 (2025). 10.1038/s41467-025-66113-z

20 Lass, A. et al. Adipose triglyceride lipase-mediated lipolysis of cellular fat stores is activated by CGI-58 and defective in Chanarin-Dorfman Syndrome. Cell Metab 3, 309–319 (2006). 10.1016/j.cmet.2006.03.005

21 Lange, M. et al. FSP1-mediated lipid droplet quality control prevents neutral lipid peroxidation and ferroptosis. Nat Cell Biol 27, 1902–1913 (2025). 10.1038/s41556-025-01790-y

22 Mao, C. et al. DHODH-mediated ferroptosis defence is a targetable vulnerability in cancer. Nature 593, 586–590 (2021). 10.1038/s41586-021-03539-7

23 Mayer, N. et al. Development of small-molecule inhibitors targeting adipose triglyceride lipase. Nat Chem Biol 9, 785–787 (2013). 10.1038/nchembio.1359

24 Althaher, A. R. An Overview of Hormone-Sensitive Lipase (HSL). ScientificWorldJournal 2022, 1964684 (2022). 10.1155/2022/1964684

25 Bisogno, T. et al. Cloning of the first sn1-DAG lipases points to the spatial and temporal regulation of endocannabinoid signaling in the brain. J Cell Biol 163, 463–468 (2003). 10.1083/jcb.200305129

26 Lin, Z. et al. The lipid flippase SLC47A1 blocks metabolic vulnerability to ferroptosis. Nat Commun 13, 7965 (2022). 10.1038/s41467-022-35707-2

27 Glomset, J. A. The mechanism of the plasma cholesterol esterification reaction: plasma fatty acid transferase. Biochim Biophys Acta 65, 128–135 (1962). 10.1016/0006-3002(62)90156-7

28 Henneberry, A. L., Wistow, G. & McMaster, C. R. Cloning, genomic organization, and characterization of a human cholinephosphotransferase. J Biol Chem 275, 29808–29815 (2000). 10.1074/jbc.M005786200

29 Sorci-Thomas, M. G., Bhat, S. & Thomas, M. J. Activation of lecithin:cholesterol acyltransferase by HDL ApoA-I central helices. Clin Lipidol 4, 113–124 (2009). 10.2217/17584299.4.1.113

30 Manthei, K. A. et al. Molecular basis for activation of lecithin:cholesterol acyltransferase by a compound that increases HDL cholesterol. Elife 7 (2018). 10.7554/eLife.41604

31 Giorgi, L. et al. Mechanistic Insights into the Activation of Lecithin-Cholesterol Acyltransferase in Therapeutic Nanodiscs Composed of Apolipoprotein A-I Mimetic Peptides and Phospholipids. Mol Pharm 19, 4135–4148 (2022). 10.1021/acs.molpharmaceut.2c00540

32 Piper, D. E. et al. The high-resolution crystal structure of human LCAT. J Lipid Res 56, 1711–1719 (2015). 10.1194/jlr.M059873

33 Horibata, Y. & Sugimoto, H. Differential contributions of choline phosphotransferases CPT1 and CEPT1 to the biosynthesis of choline phospholipids. J Lipid Res 62, 100100 (2021). 10.1016/j.jlr.2021.100100

34 Plutner, H., Davidson, H. W., Saraste, J. & Balch, W. E. Morphological analysis of protein transport from the ER to Golgi membranes in digitonin-permeabilized cells: role of the P58 containing compartment. J Cell Biol 119, 1097–1116 (1992). 10.1083/jcb.119.5.1097

35 Ibrahim, I. M., Abdelmalek, D. H. & Elfiky, A. A. GRP78: A cell’s response to stress. Life Sci 226, 156–163 (2019). 10.1016/j.lfs.2019.04.022

36 Nakamura, N., Lowe, M., Levine, T. P., Rabouille, C. & Warren, G. The vesicle docking protein p115 binds GM130, a cis-Golgi matrix protein, in a mitotically regulated manner. Cell 89, 445–455 (1997). 10.1016/s0092-8674(00)80225-1

37 Lu, L. & Hong, W. Interaction of Arl1-GTP with GRIP domains recruits autoantigens Golgin-97 and Golgin-245/p230 onto the Golgi. Mol Biol Cell 14, 3767–3781 (2003). 10.1091/mbc.e03-01-0864

38 Lujan, P. et al. Sorting of secretory proteins at the trans-Golgi network by human TGN46. Elife 12 (2024). 10.7554/eLife.91708

39 Henneberry, A. L., Wright, M. M. & McMaster, C. R. The major sites of cellular phospholipid synthesis and molecular determinants of Fatty Acid and lipid head group specificity. Mol Biol Cell 13, 3148–3161 (2002). 10.1091/mbc.01-11-0540

40 Lei, G., Zhuang, L. & Gan, B. Targeting ferroptosis as a vulnerability in cancer. Nat Rev Cancer 22, 381–396 (2022). 10.1038/s41568-022-00459-0

41 Sehm, T. et al. Sulfasalazine impacts on ferroptotic cell death and alleviates the tumor microenvironment and glioma-induced brain edema. Oncotarget 7, 36021–36033 (2016). 10.18632/oncotarget.8651

42 Tao, L. et al. Integrative clinical and preclinical studies identify FerroTerminator1 as a potent therapeutic drug for MASH. Cell Metab 36, 2190–2206 e2195 (2024). 10.1016/j.cmet.2024.07.013

43 Dowdy, S. F. Overcoming cellular barriers for RNA therapeutics. Nat Biotechnol 35, 222–229 (2017). 10.1038/nbt.3802

44 Boland, M. L. et al. Towards a standard diet-induced and biopsy-confirmed mouse model of non-alcoholic steatohepatitis: Impact of dietary fat source. World J Gastroenterol 25, 4904–4920 (2019). 10.3748/wjg.v25.i33.4904

45 Pratt, D. S. & Kaplan, M. M. Evaluation of abnormal liver-enzyme results in asymptomatic patients. N Engl J Med 342, 1266–1271 (2000). 10.1056/NEJM200004273421707

46 Forte, T. M. et al. Targeted disruption of the murine lecithin:cholesterol acyltransferase gene is associated with reductions in plasma paraoxonase and platelet-activating factor acetylhydrolase activities but not in apolipoprotein J concentration. Journal of Lipid Research 40, 1276–1283 (1999). 10.1016/s0022-2275(20)33489-1

47 Guo, M. et al. Spontaneous Atherosclerosis in Aged LCAT-Deficient Hamsters With Enhanced Oxidative Stress-Brief Report. Arterioscler Thromb Vasc Biol 40, 2829–2836 (2020). 10.1161/ATVBAHA.120.315265

48 Chu, Q. et al. Repurposing a tricyclic antidepressant in tumor and metabolism disease treatment through fatty acid uptake inhibition. J Exp Med 220 (2023). 10.1084/jem.20221316

49 Zhai, X. et al. AMPK-regulated glycerol excretion maintains metabolic crosstalk between reductive and energetic stress. Nat Cell Biol 27, 141–153 (2025). 10.1038/s41556-024-01549-x

