## Supplementary Figures for "CHPT1–LCAT rewires lipolysis towards ferroptosis"

**Extended Data Fig.1 | MAGL inhibitors promote ferroptosis**

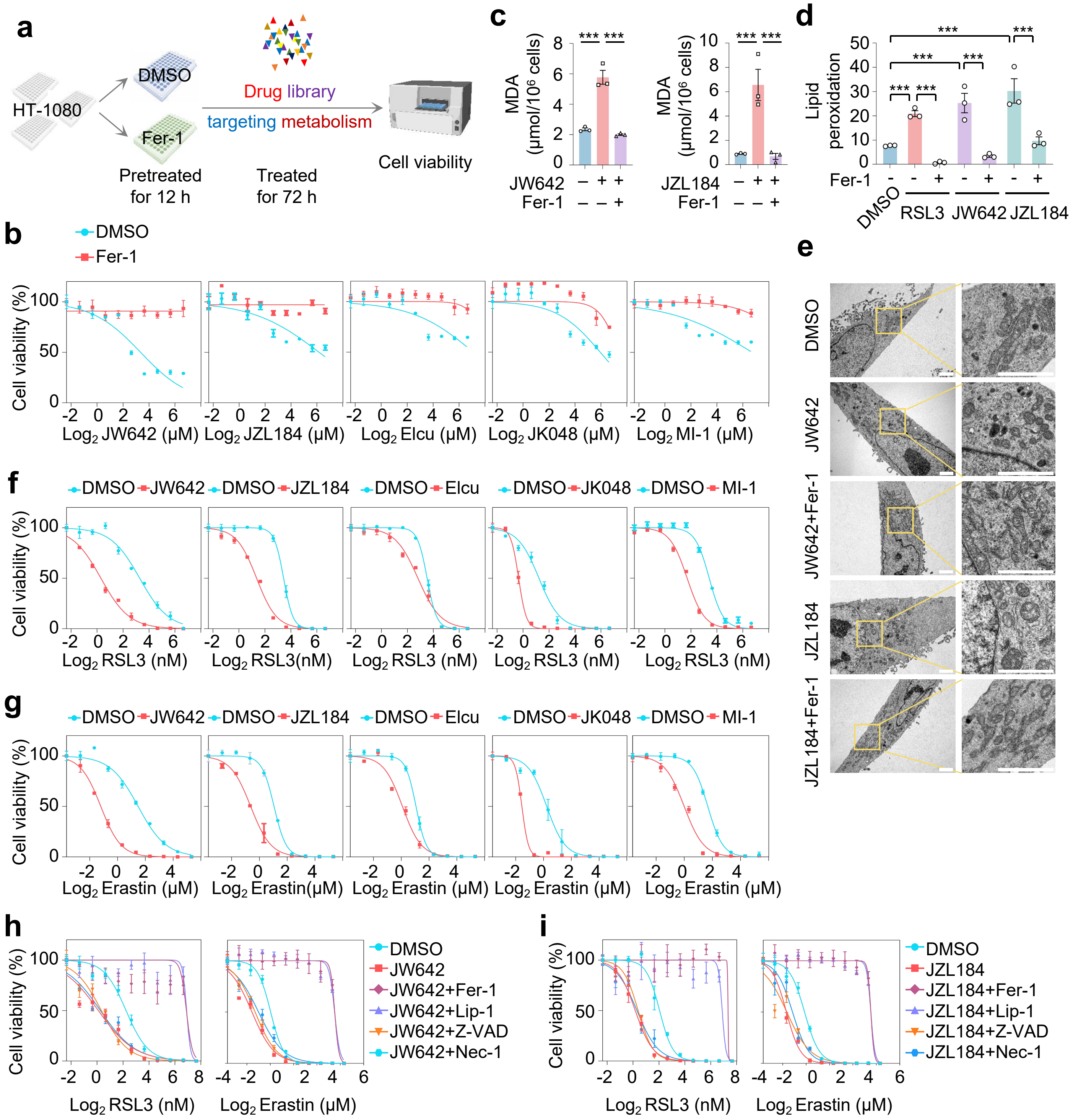

### Extended Data Fig.2 | ATGL Inhibition attenuates ferroptosis

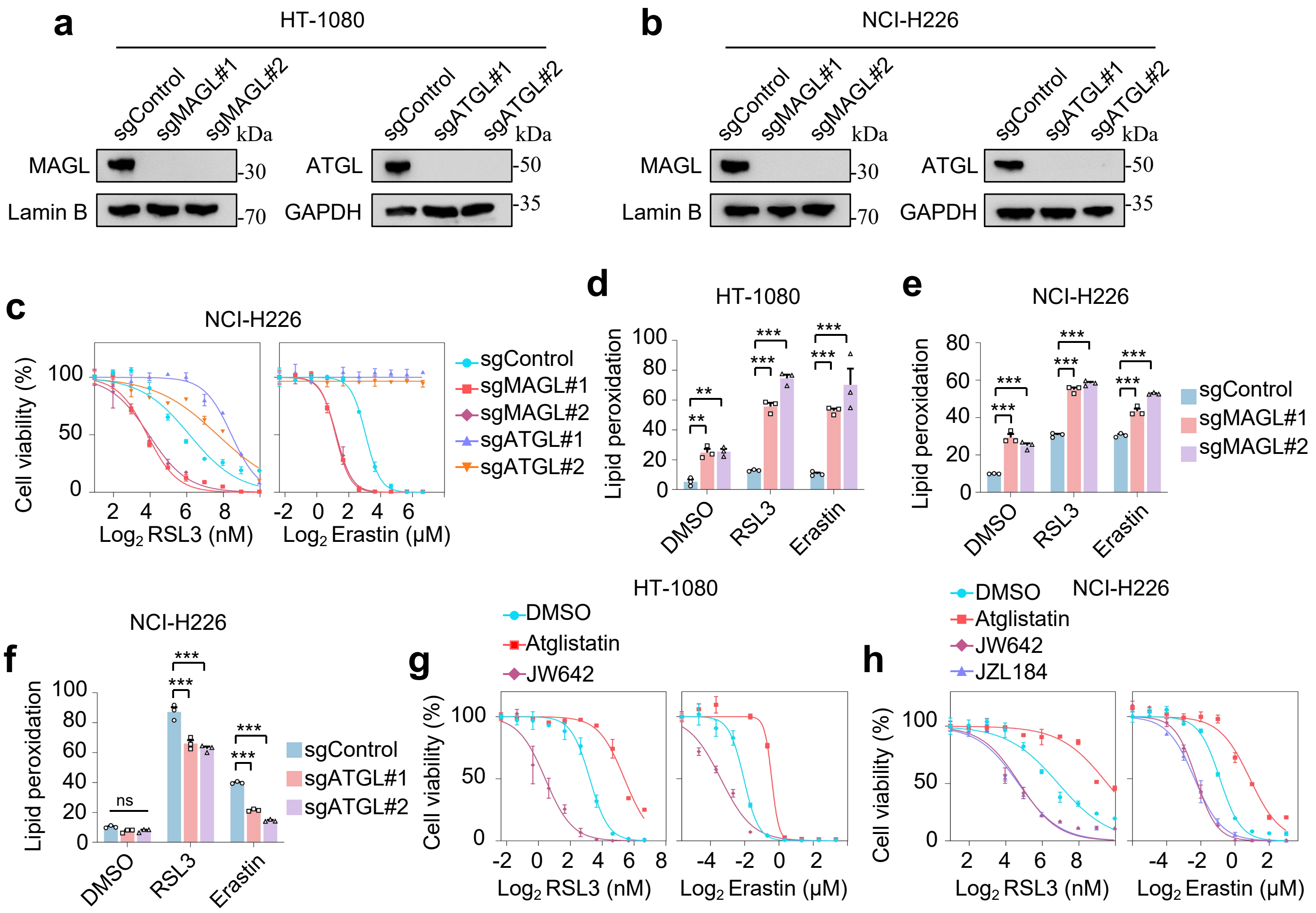

**Extended Data Fig.3 | DAGL inhibitors sensitize cancer cells to ferroptosis**

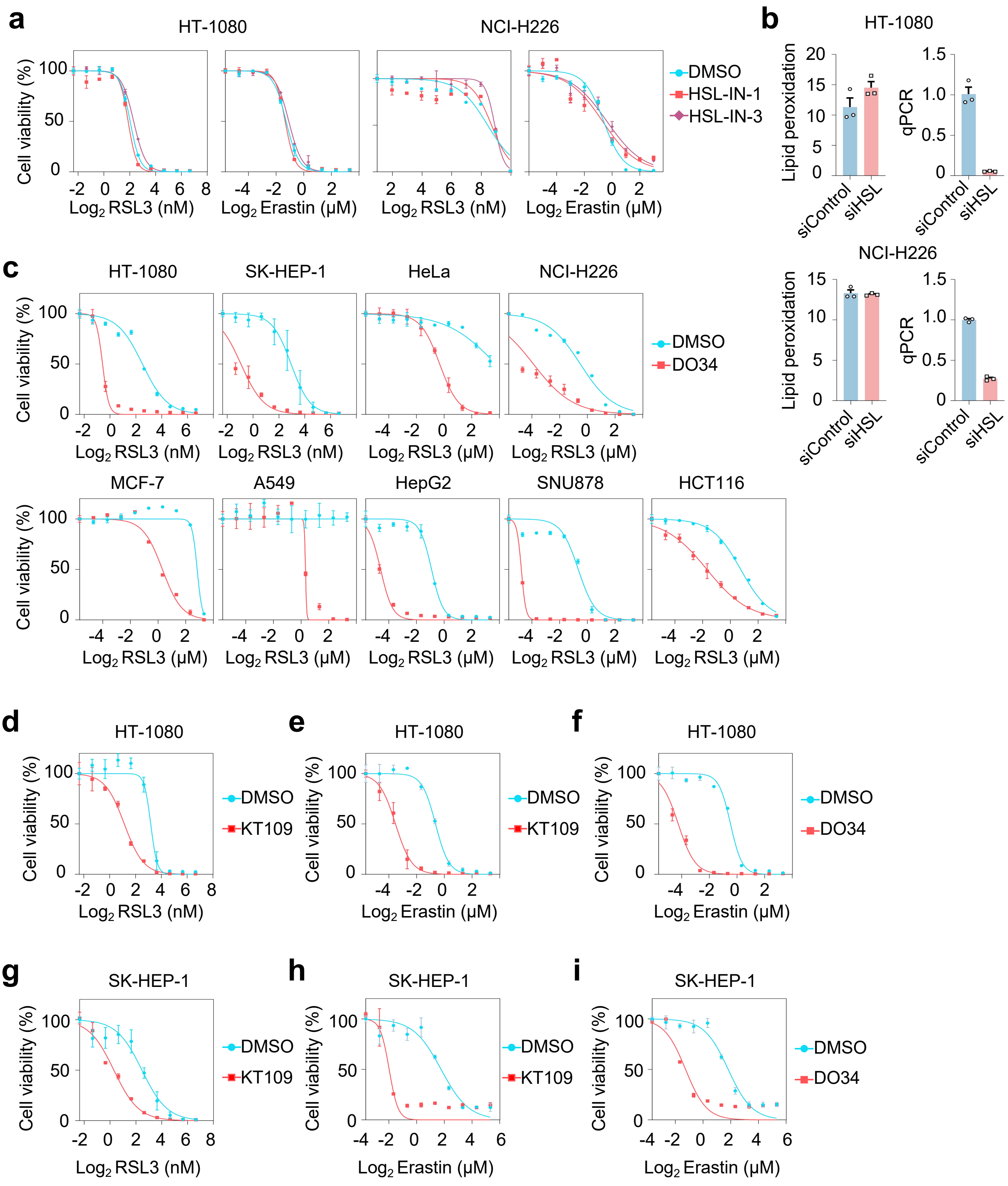

**Extended Data Fig.4 | DAG determines the sensitization of cancer cells to ferroptosis**

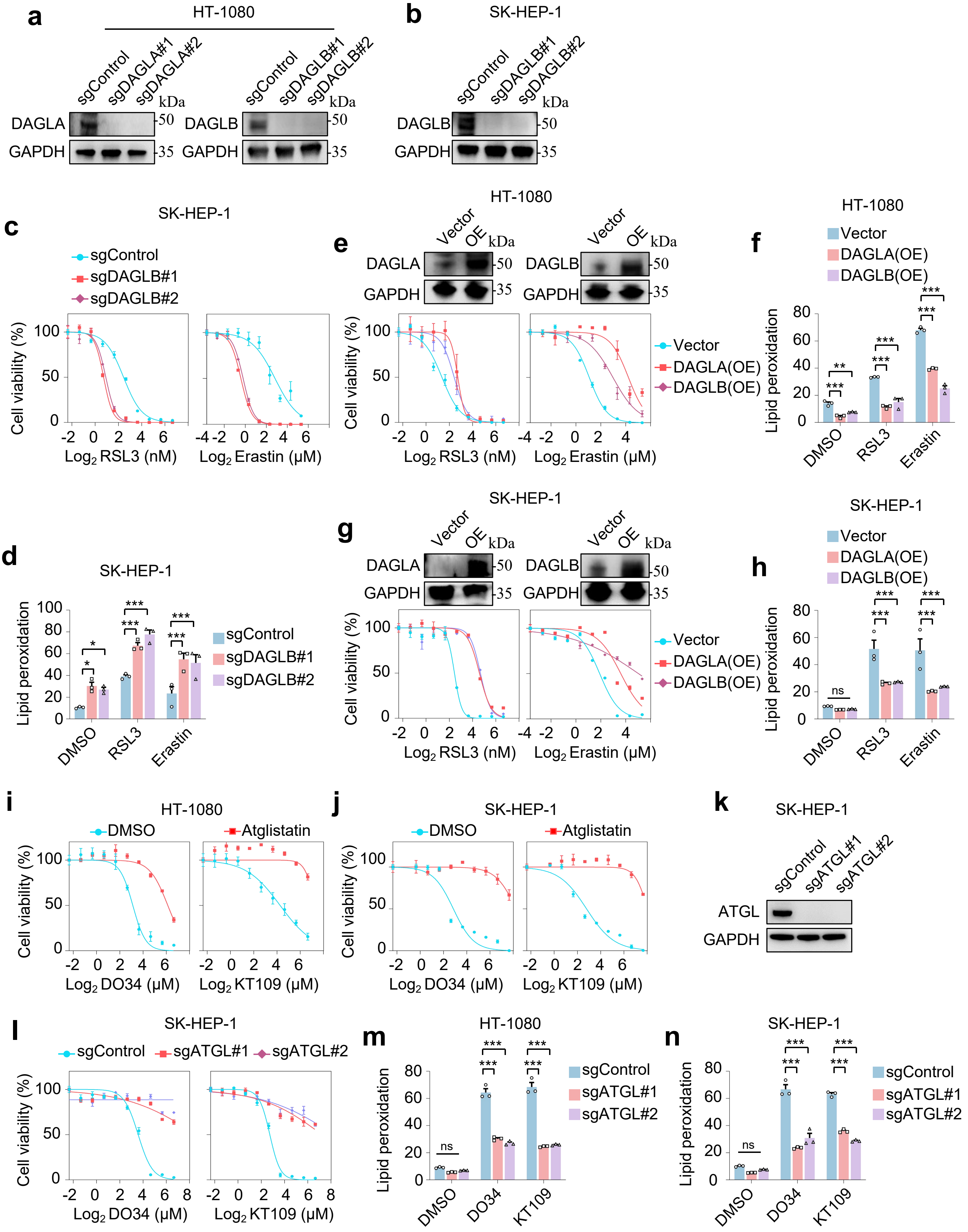

**Extended Data Fig.5 | DAGL inhibition induces ferroptosis**

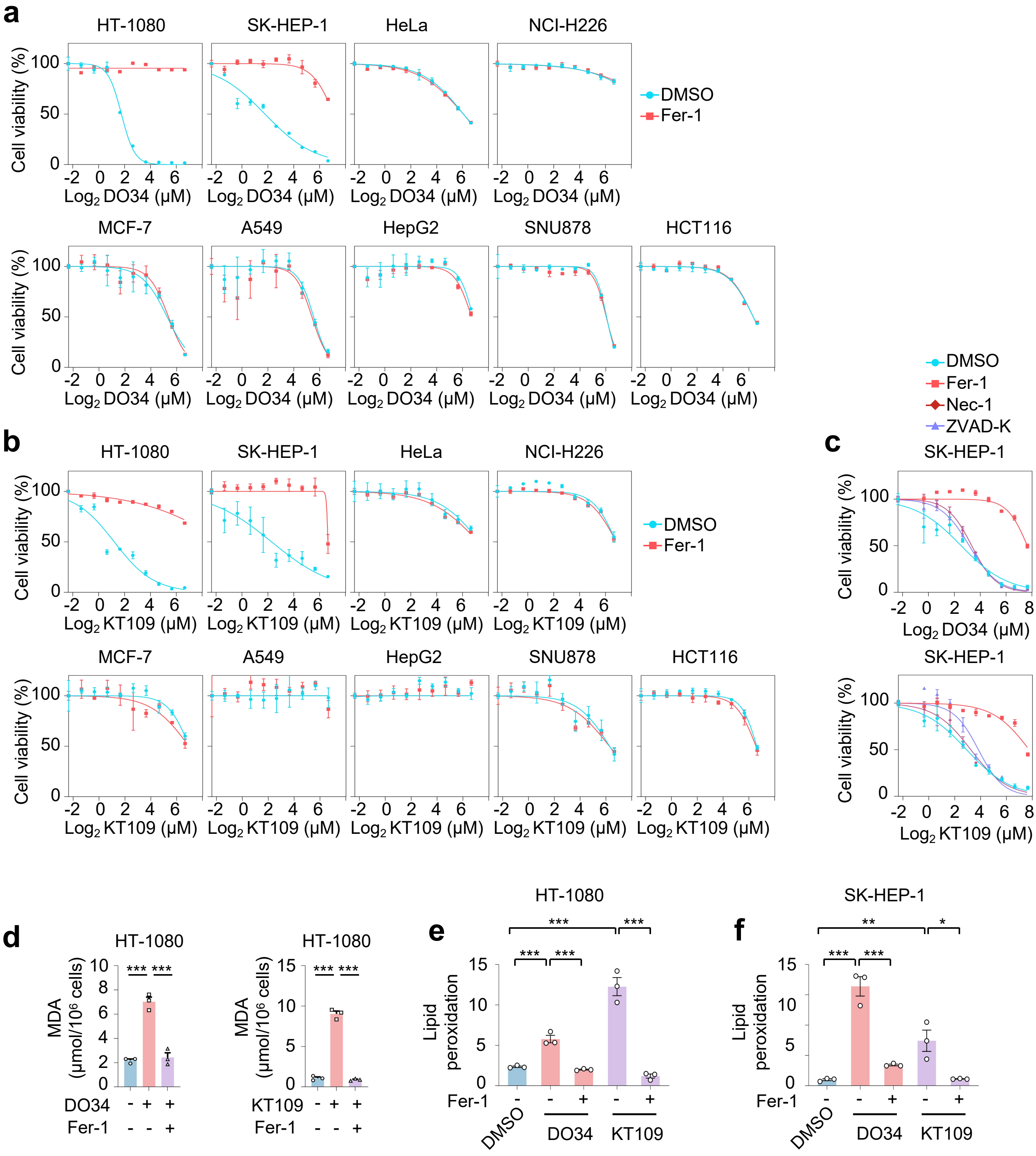

**Extended Data Fig.6 | Cholesteryl ester accumulation is a causal determinant of ferroptosis**

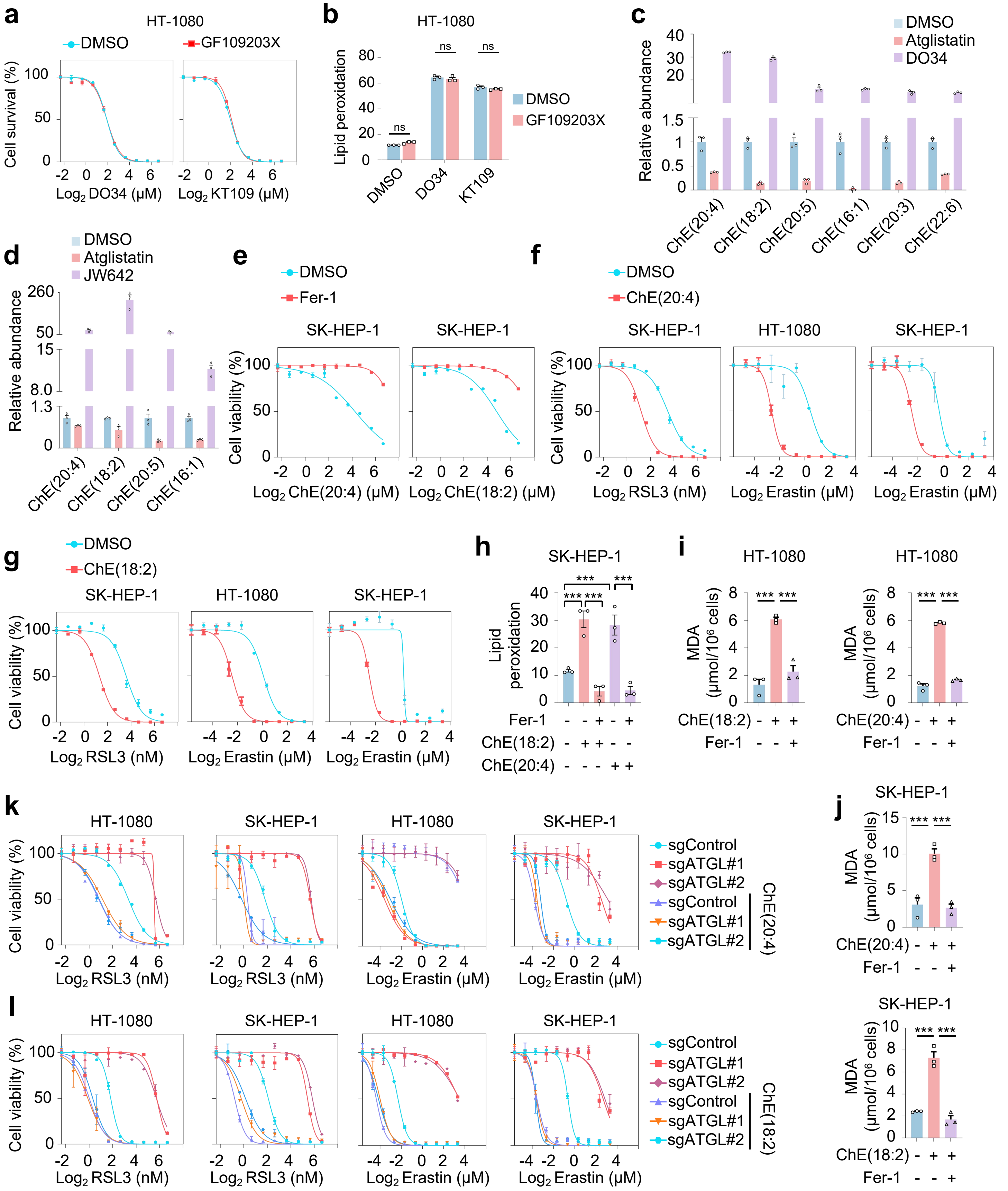

**Extended Data Fig.7 | siRNA screening identifies CHPT1 and LCAT as mediators of DAGL inhibition-induced ferroptosis**

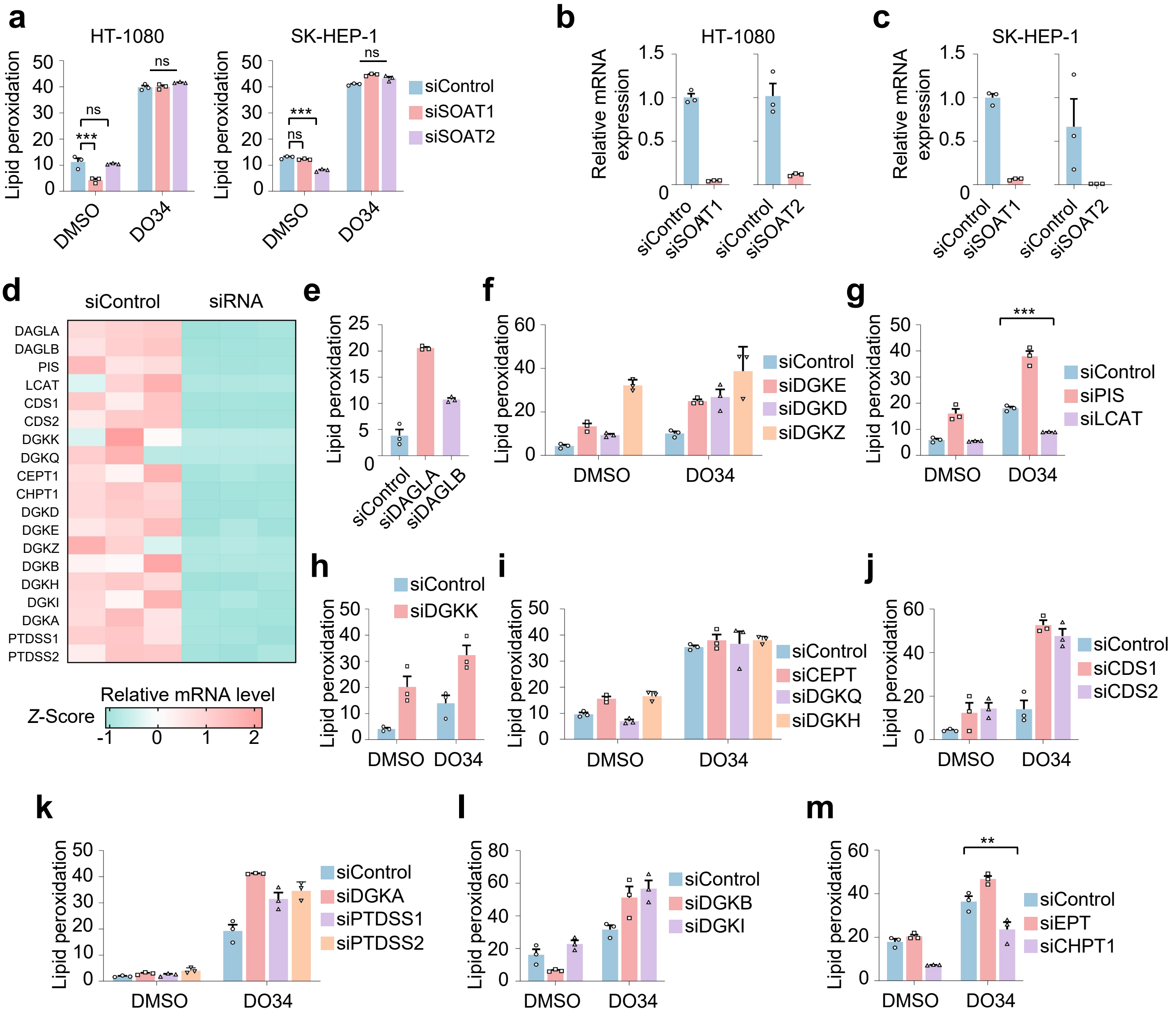

**Extended Data Fig.8 | The CHPT1–LCAT axis regulates DAG routing to ferroptosis**

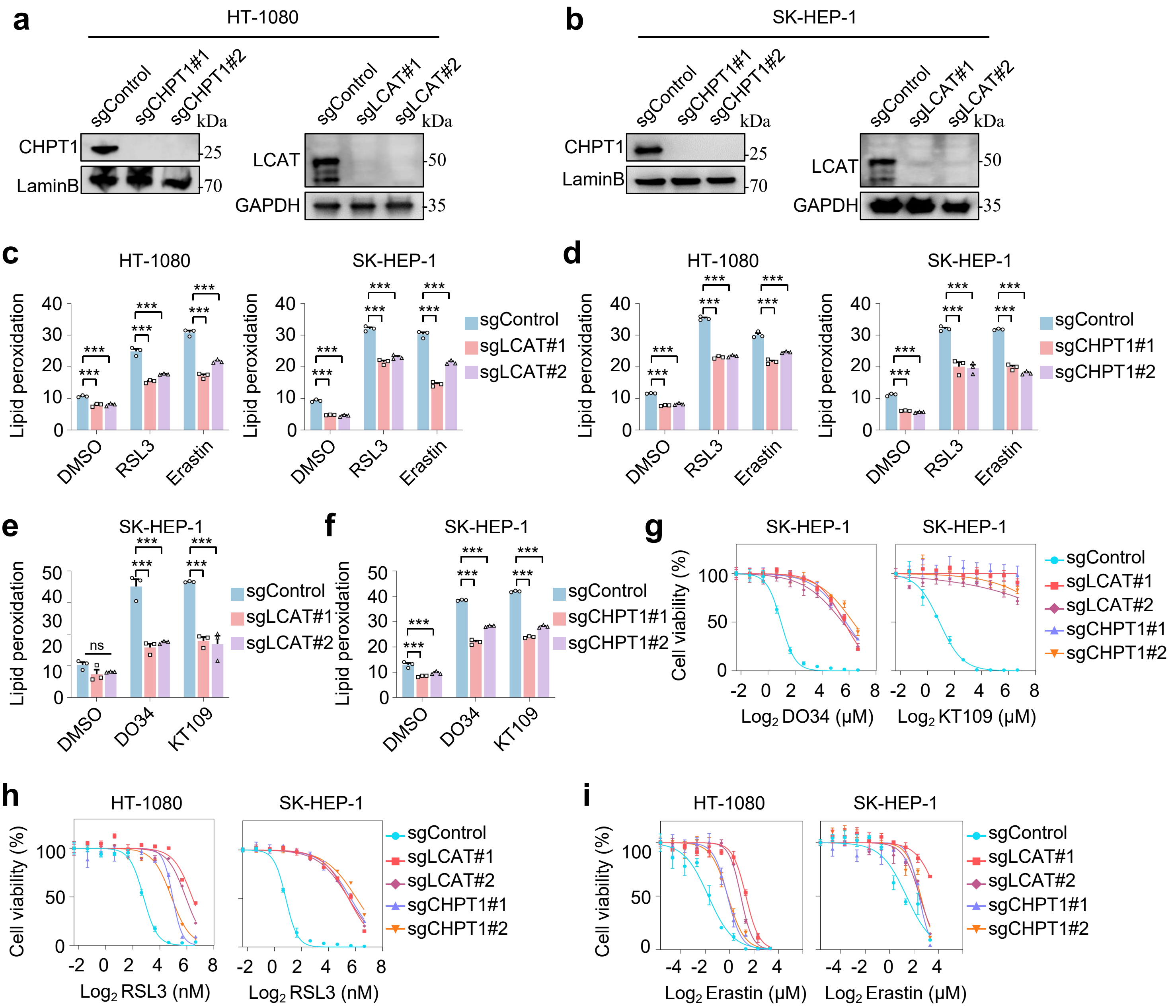

**Extended Data Fig.9 | Intracellular LCAT mediates ferroptosis**

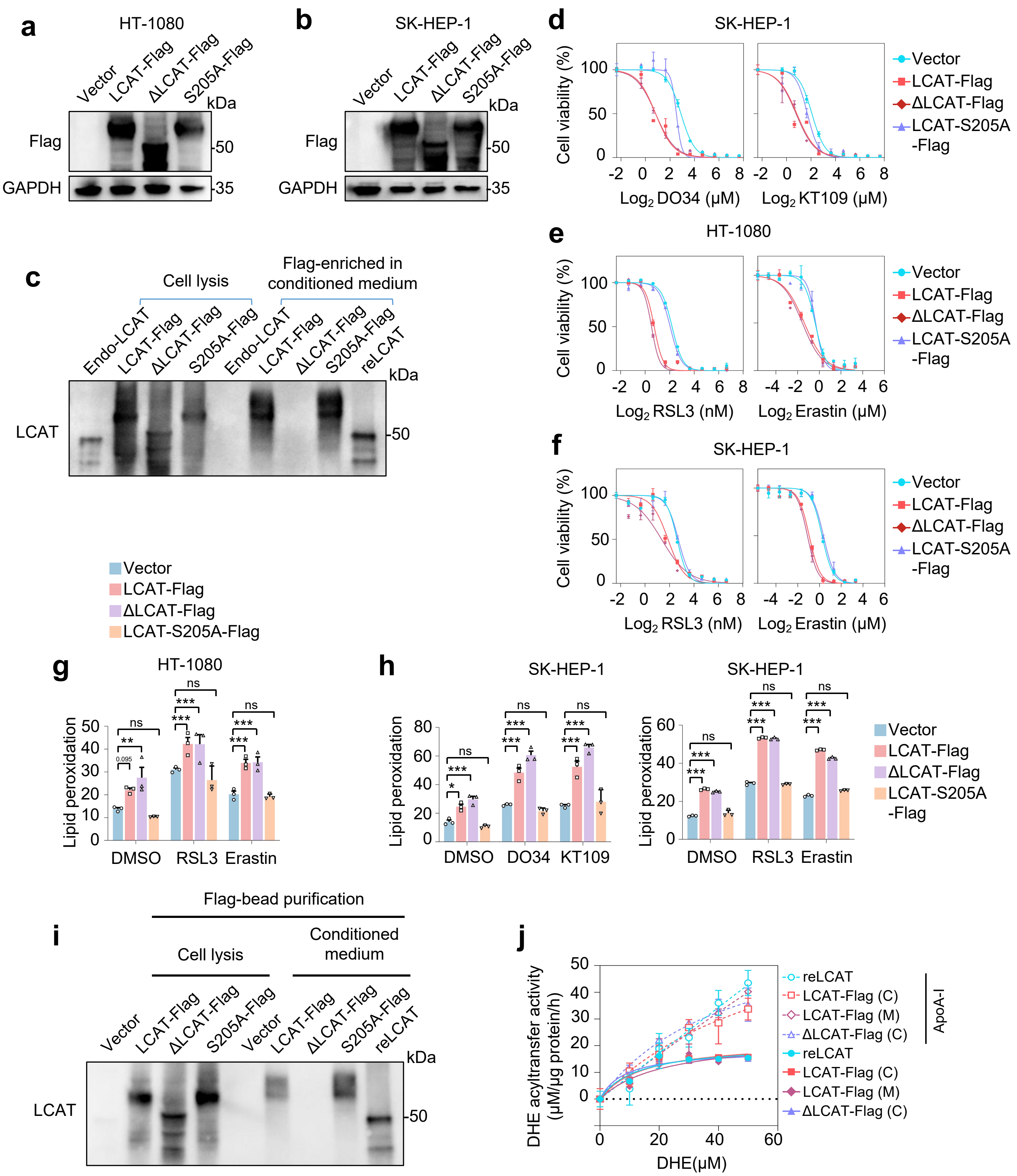

**Extended Data Fig.10 | Intracellular localization of CHPT1 and LCAT**

a

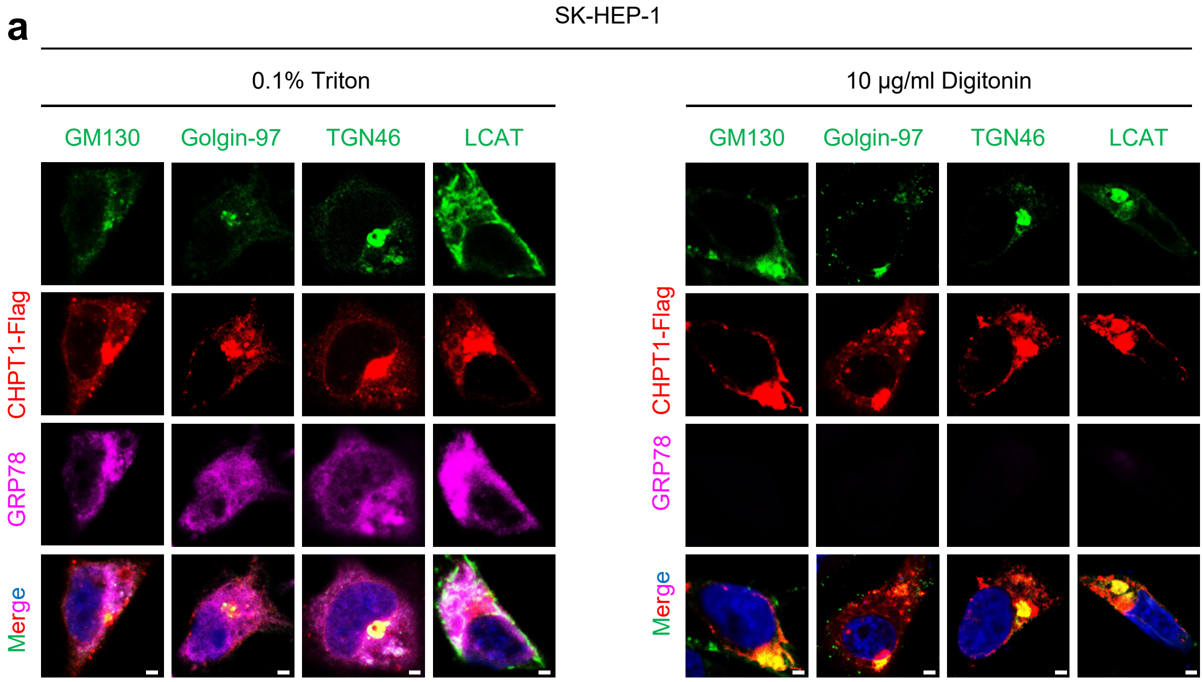


b

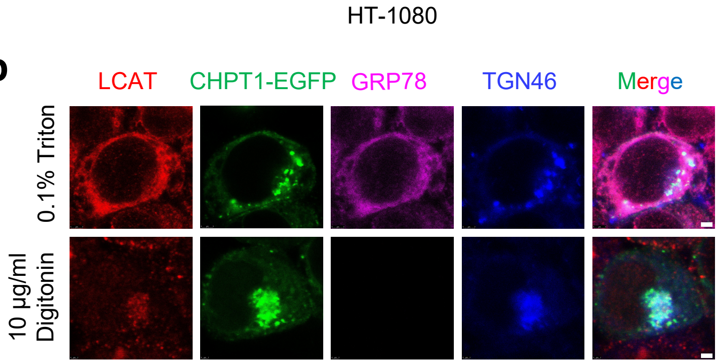

**Extended Data Fig.11 | CHPT1–LCAT mediates ferroptosis across diverse contexts**

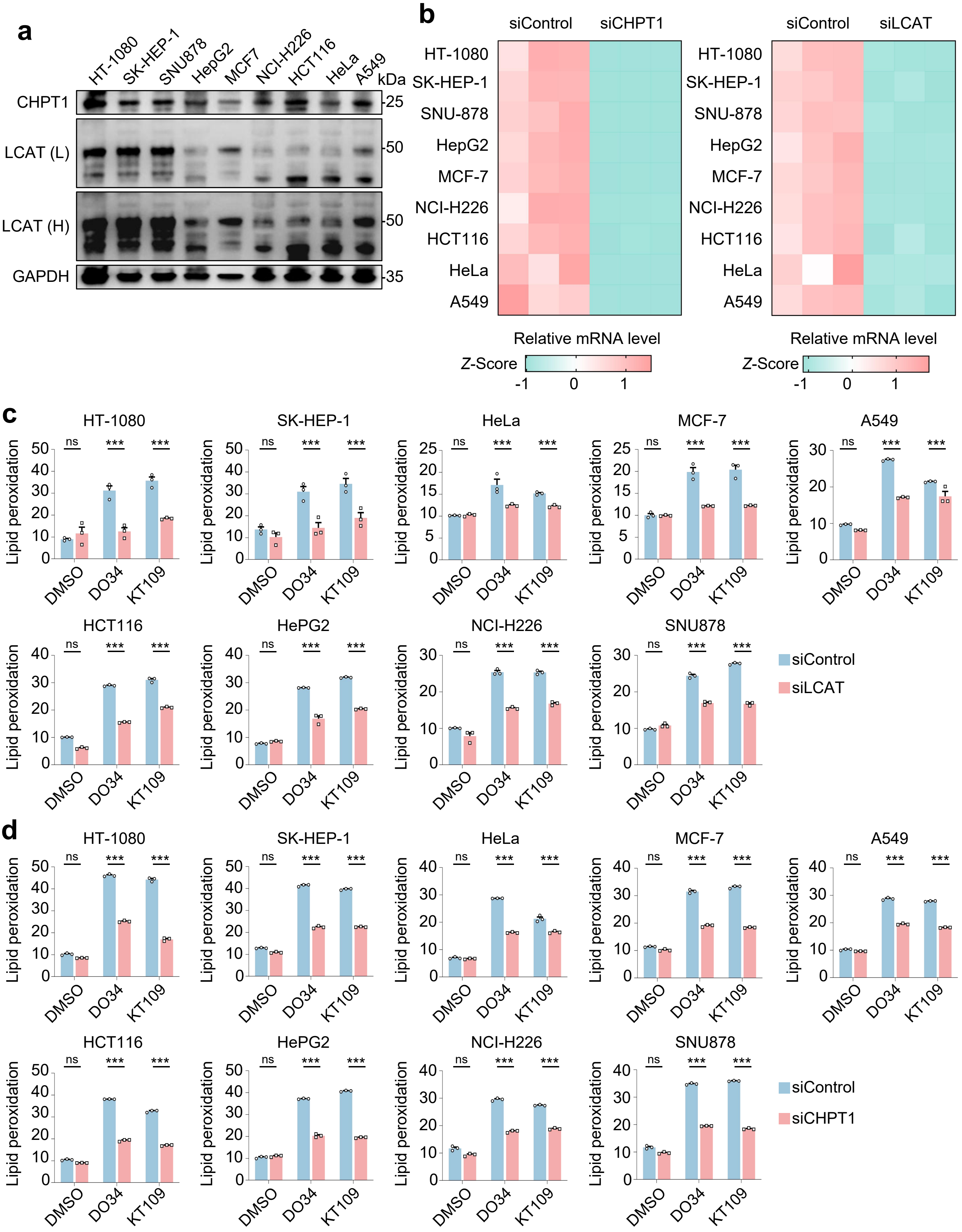

**Extended Data Fig.12 | MAGL and DAGL inhibitors suppress tumour growth via ferroptosis**

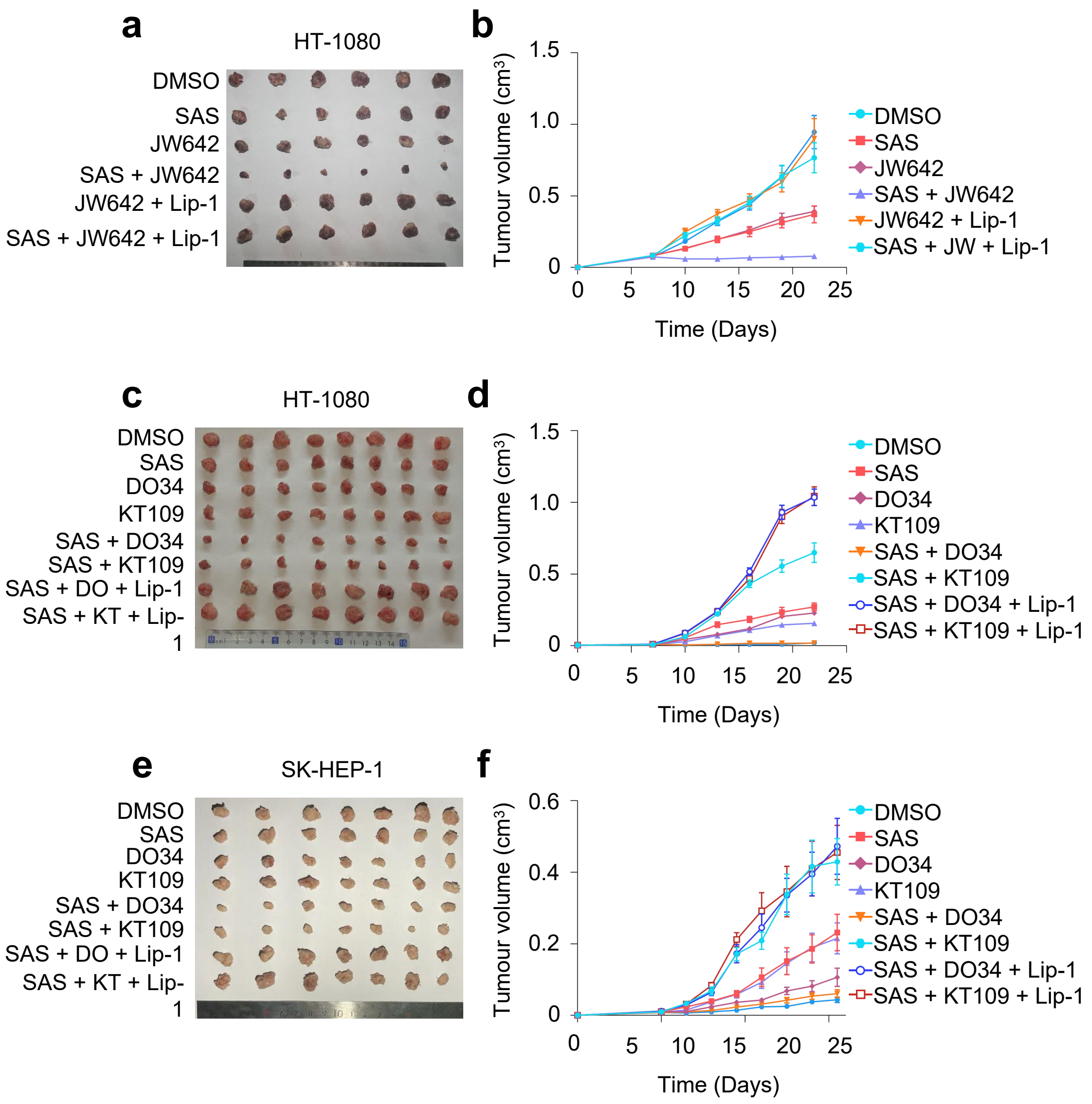

**Extended Data Fig.13 | GalNAc–siRNA enables hepatic knockdown of CHPT1 and LCAT in mice**

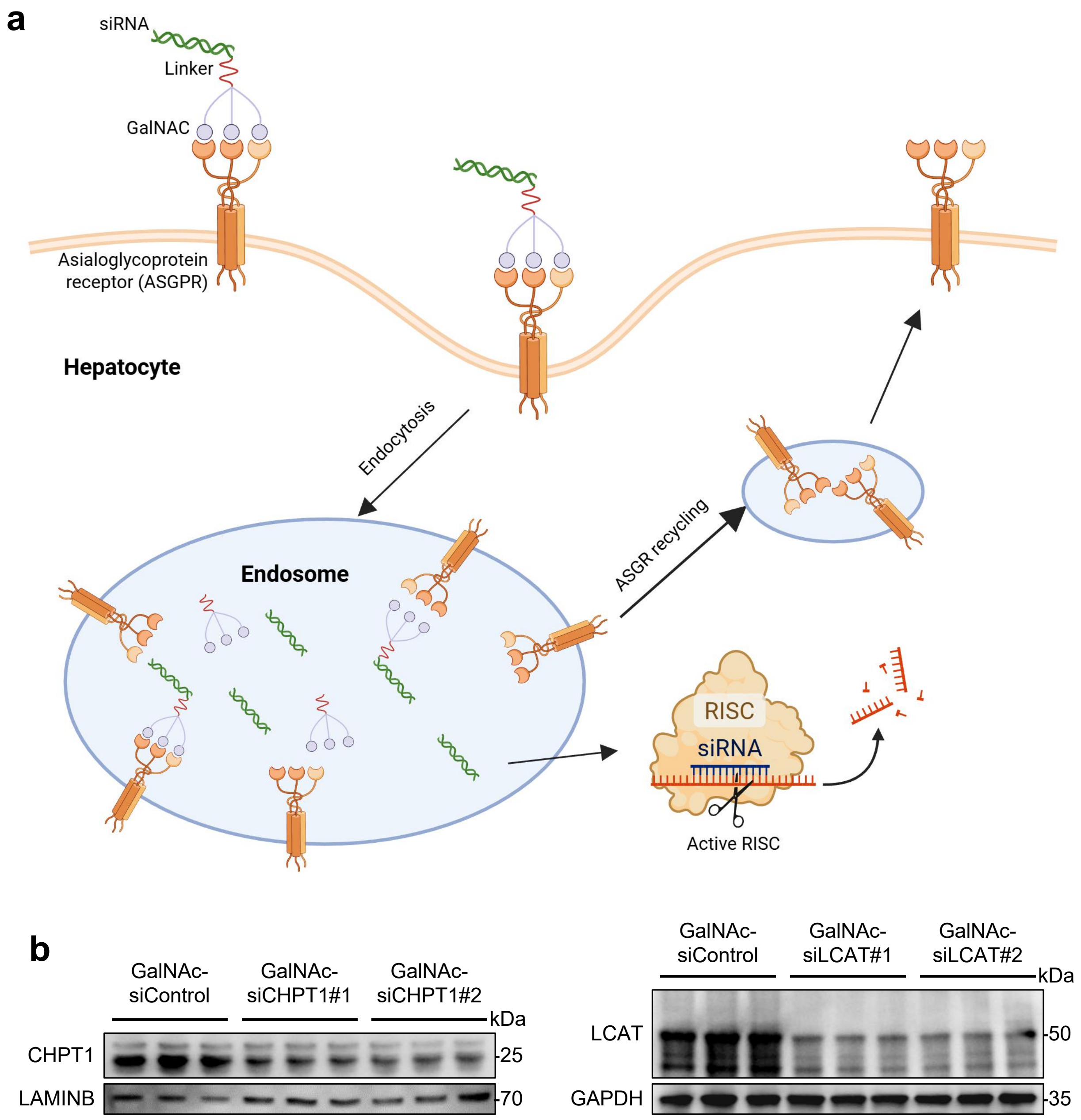

**Extended Data Fig.14 | Targeting the CHPT1–LCAT axis attenuates MASH in mice**

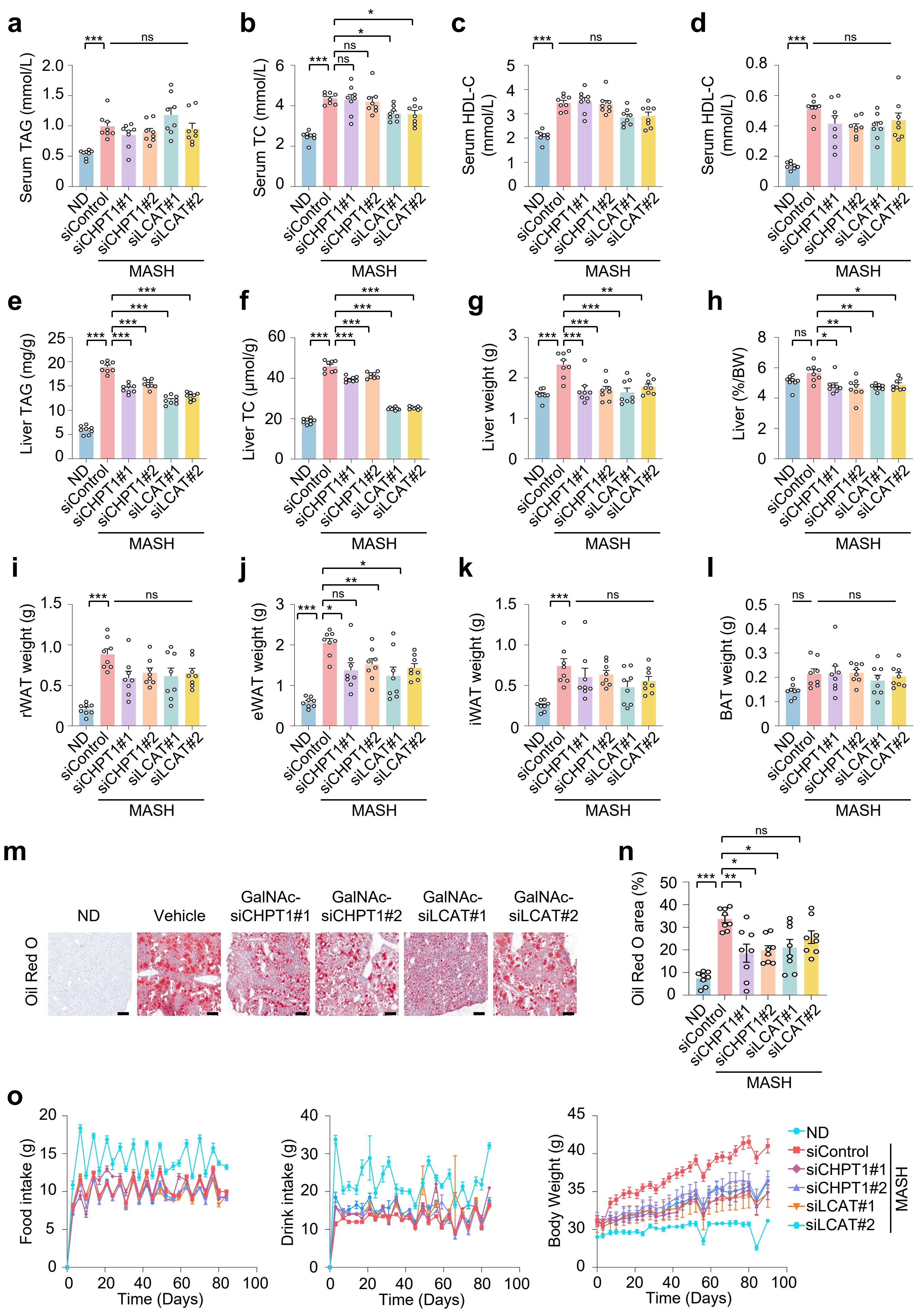

**Extended Data Fig.15 | DAG routing drives ferroptosis through a CHPT1–LCAT metabolic axis**

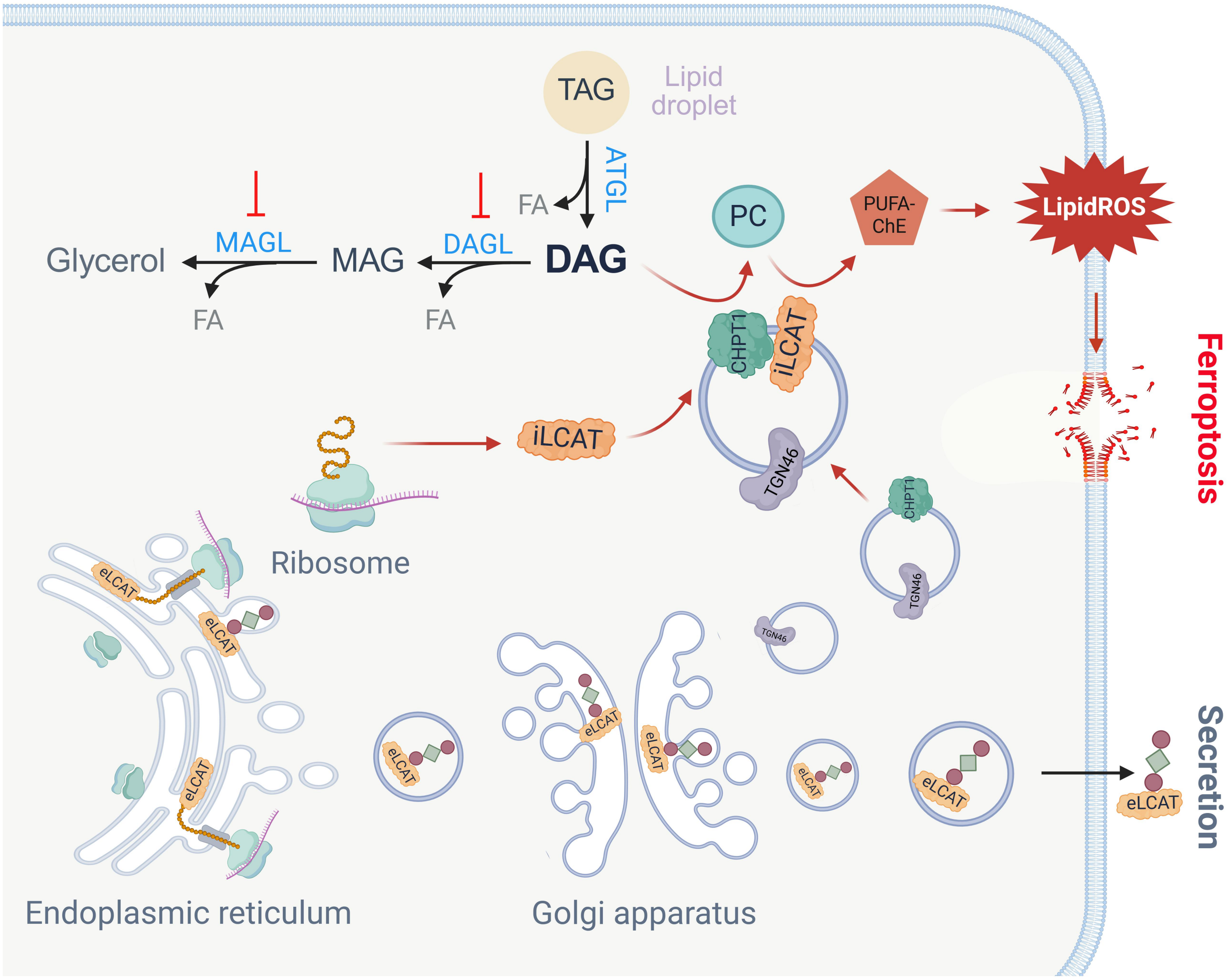
